# Conditional Dirichlet Flow Matching for exploring the sequence space of protein families

**DOI:** 10.64898/2026.11.26.769518

**Authors:** Sebastian Andersson, Nina J. Schuch, Ingemar André

## Abstract

Protein folds have evolved to support a diverse range of functions, and repurposing an existing fold can offer a more effective route to engineering new functions than designing a scaffold *de novo*. Here, we introduce Syzygy, a generative model that learns the evolutionary sequence patterns of protein families to generate artificial homologues that diverge from natural sequences while preserving family-specific characteristics. Based on Dirichlet flow matching, Syzygy is trained on a single multiple sequence alignment and enables sequence generation conditioned on protein length, a discrete functional label, or both. The model is computationally efficient, with average training times of under 10 minutes on a single GPU and generation of 1,000 sequences in approximately 30 seconds. Evaluation across four protein families shows that generated sequences are predicted to retain their respective family folds with high structural confidence. When conditioned on fluorescence colour within the green fluorescent protein (GFP) family, 98% of generated sequences match the requested sequence-based colour classification, including blue fluorescent proteins, which constitute only 2.4% of the training data. Experimental characterisation of selected GFP and NDM-1 designs demonstrates that artificial homologues can retain fluorescence and enzymatic activity, although expression levels remain lower than those of their wild-type counterparts. Syzygy thus provides a rapid and controllable approach to generating synthetic protein families and functional starting scaffolds for further protein engineering, enabling exploration of regions of family-specific sequence space not represented among known natural homologues.

## 1. Introduction

Many protein engineering tasks do not need unrestricted *de novo* generation. They require a way to explore sequence diversity inside a known family while preserving family-specific structural and functional constraints. Such a tool should operate on sequence alone, explore the natural length variation in a family, and enable control over properties of the generated sequences. While recent advances in structure-based generative models have revolutionised *de novo* design and enabled a straightforward structure-to-sequence workflow with high success rates[1, 2], they do not meet these criteria due to their reliance on structural priors. Designing artificial homologues with these tools requires added constraints, such as PSSM-based biases or fixing of amino acid identities at specific positions, because a backbone alone does not preserve family-specific residues[3]. Furthermore, these pipelines are typically computationally intensive and do not explore length variation in the generative model.

Sequence-only models of protein families offer a compelling alternative, and have a long history. Potts models[4], variational autoencoders[5], generative adversarial networks[6], and latent diffusion models[7] have been shown to result in experimentally validated proteins, with autoregressive Potts-type models having substantially training times than Boltzmann-machine DCAs[8]. These models take the family alignment as their main input. Some models also take an addition label: a three-class solubility label in a luciferase-family autoencoder[5], or a fitness label in a latent diffusion model tested on GFP[7].

More recently, sequence-only modelling has been dominated by large models trained across protein space, using autoregressive or discrete diffusion frameworks[9–11]. They offer a wide range of control: a family can be specified by a prompt of homologous sequences[12], or by control tags and fine-tuning[9]. These models are the natural choice when the family is not known in advance or when few sequences are available. They have hundreds of millions to tens of billions of parameters and are trained for days on a vast dataset of protein sequences. These models are built to capture broad evolutionary patterns across the whole of protein space, and their scale and generality are central to that goal. This breadth, however, comes with trade-offs for tasks scoped to a single protein family: conditioning is typically supported only at the family or organism level, so steering generation toward a specific function or property within a family generally requires additional fine-tuning. The computational resources and inference time required to model the full diversity of protein space likewise exceed those needed for a single-family task, and rare variants within a specific local fitness landscape can be underrepresented without further adaptation.

We introduce a complementary paradigm: a lightweight, task-specific generative model for controllable protein sequence design where both sequence length and a discrete functional class can be specified, in one network that generates all positions in parallel. Dirichlet flow matching, introduced for DNA sequences[13], is a suitable basis: it models categorical data on the probability simplex, generates all positions in parallel, and trains with a simple regression objective. Here we train a small Dirichlet flow matching model, Syzygy, for each family from one multiple sequence alignment (MSA), adapting the model to protein sequences. We replace the convolutional network of the original formulation with a Transformer conditioned through AdaLN-Zero[14], and use this pathway to pass the sequence length, a class label, or both. A sampling parameter, *α*_max_, sets the balance between similarity to the original family and sequence diversity. The models have 5.0 to 5.3 million parameters, train in under 10 minutes on average on one A100 GPU, and generate 1000 sequences in about 30 seconds. Sampling needs no structure, scoring matrix, or pretrained model.

We train the model on four families that differ in alignment depth, length, and function: three enzymes and one fluorescent protein. We show that the generated sequences keep the family fold, are confidently predicted, are consistent with the training sequences, and that conditioning inputs are able to reliably control the output. Label conditioning has equal success rates for both over-represented and under-represented classes. Length conditioning follows the target within the main length range of the training alignment. We tested 8 GFP designs in the lab, all with less than 95% identity to any training sequence: green and blue designs fluoresce as designed, while yellow and cyan show green fluorescence. For NDM-1, we tested 8 designs: two designs displayed activity at approximately a third of the level of the wild type without additional optimisation. Syzygy provides a fast and controllable alternative for family-specific protein design and complements existing structure-based design workflows.

## 2. Results

### 2.1. Overview of the framework

We developed Syzygy, a generative model for family-conditioned protein sequence design based on Dirichlet flow matching, an extension of continuous normalising flows to data on a probability simplex[13]. The model learns a vector field that transports an uninformative Dirichlet prior toward the amino-acid distribution characteristic of a target protein family, learned directly from a family-specific multiple sequence alignment. We adapted this framework to the 21-character amino acid alphabet (20 standard residues plus a gap character) and replaced the convolutional architecture of the original formulation with a Transformer parameterised by adaptive layer normalisation (AdaLN-Zero)[14], enabling the network to condition generation on auxiliary properties (sequence length, a discrete functional label, or both together) in addition to the diffusion time variable itself (full architectural details can be seen in figure 9).

Dirichlet flow matching was originally developed for DNA sequence design[13]; we extend it here to protein sequences. Unlike a naive linear interpolation on the simplex, which becomes discontinuous and increasingly sensitive to integration step size as the alphabet size grows, Dirichlet flow matching remains well-behaved for the 21-character amino acid alphabet, making it a more robust choice for categorical sequence data of this kind.

Sequences are generated by numerically integrating the learned vector field toward a chosen terminal concentration parameter, *α*_max_, which controls how peaked, and therefore how confident, the final per-position amino-acid distribution is: higher values yield more deterministic, training-set-like sequences; lower values retain more diversity. In order to characterise the method, we trained Syzygy independently on four protein families chosen to span differing alignment depth, sequence length, and functional diversity.

### 2.2. Protein families and MSA curation

We generated multiple sequence alignments for four protein families: protein tyrosine phosphatases (PTPs)[15], Flavin oxidoreductases (FREs)[16], New Delhi Metallo-*β*-lactamase (NDM-1)[17], and green fluorescent proteins (GFPs)[18, 19].

We selected these protein families because their catalytic residues and substrate specificity have been experimentally determined. For three of the four families, experimentally determined structures were also available[20–22]. At the time of this study, no experimentally determined structures were available for the selected FRE model protein. However, experimental studies have identified residues that are important for catalysis in the enzyme families or for fluorescence in the GFPs. We used those experimentally characterised residues as a reference for evaluating whether our sequence-based model could learn biologically relevant features without incorporating structural or functional information. The experimentally characterised residues used as references are summarised in Table S4.

The PTP alignment represents proteins that have a key role in cellular signal transduction by removing phosphate groups from tyrosine residues. The catalytic domain holds the active site, which is formed across several structured loops. This catalytic machinery is highly conserved throughout the family and defined by several loops that contain key residues for phosphorylation. PTP1 is a highly dynamic enzyme in which the catalysis is regulated by allosteric interactions and conformational motions[15, 23].

The FRE alignment was based on the flavin oxidoreductase from *Vibrio harveyi* that catalyses the reduction of flavin using NADPH as a cofactor[24]. The enzyme preferentially uses riboflavin as an electron acceptor. No structure has been solved in complex with substrate. However, the crystal structure of a homologous FRE protein from *E*.*coli* provides structural insight into the isoalloxazine-binding site, which is characterised by hydrophobic interactions with the flavin. Several conserved residues have also been identified as important for flavin binding. In contrast, the NADPH-binding site is less well characterised, although structural models have been proposed, placing it in close proximity to the flavin-binding site, which facilitates efficient electron transfer. Given the relatively high sequence identity between the homologous *E*.*coli*[25] FRE and the selected FRE protein (48%), these structural features provide a basis for proposing that the flavin-binding site and its associated conserved residues are similarly organised in the selected protein.

NDM-1 is a metallo-*β*-lactamase that requires two zinc ions for its catalytic activity[26]. The enzyme hydrolyses *β*-lactam antibiotics, especially carbapenems, which are the most specific substrates for NDM-1. Its active site is formed by a set of conserved metal/coordinating residues together with several flexible loops surrounding the catalytic site. Seven residues contribute to the active site, and these have been used as reference residues in this study. In addition, length variation in the N-terminal region of NDM-1 was considered in the analysis because previous work has shown that expression of NDM-1 can be increased by removing part of the N-terminus. In particular, truncating the first 38 residues has been shown to maximise protein yields[27].

The GFP dataset represents a highly conserved family of fluorescent proteins with a characteristic *β–*barrel surrounding the chromophore. GFP variants have been widely studied and extensively used as a molecular imaging tool. The fluorescence comes from an autocatalytically formed chromophore built by residues inside the barrel. The properties of fluorescence in the GFP family are determined by the structure and the electronic environment of the chromophore. Therefore, the amino acid residues around the chromophore are very conserved, as they influence the protonation state, hydrogen bonding, electrostatic interactions and conformational state, which can alter the observed excitation and emission maxima. In addition, mutations that affect protein folding and conformational dynamics may influence chromophore maturation, resulting in changes in fluorescence intensity and maxima[28–31]

#### 2.2.1. Effect of MSA depth and sequence composition on prediction

The initial prediction of new sequences was based on MSAs that have been highly filtered to reduce gap percentages. However, this procedure reduced the MSA depth to only 200 sequences for the NDM-1 family. Compared to the other studied families, NDM-1 showed worse quality in evaluation metrics. Increasing the MSA depth from 200 to 400 and then to 800 sequences resulted in progressively improved computational results in this system.

However, increasing the number of sequences was not universally beneficial. This could particularly be seen for the GFP family, for which the initial alignment contained a large number of sequences. The alignment for GFP also contained many red fluorescent proteins (RFPs). We found that their inclusion substantially reduced the quality of generated sequences. While GFPs and RFPs share a common evolutionary origin, they substantially differ in chromophore composition. Removing the RFP sequences improved sequence quality, especially for Blue Fluorescent Proteins (BFPs), which are the least represented fluorophore class in the MSA.

All four MSAs have comparable sequence depth, but differ in the degree of variation in length and identity between sequences (Table 1). Even though the gap content is relatively similar between the alignments, ranging from 57.65% for NDM-1 to 65.15% for PTP1, the interdecile range (IDR) of the protein length distributions differed substantially between families. The IDR, which captures the spread of the middle 80% of protein lengths, therefore indicates variation in sequence length across the families.

**Table 1.** Protein family composition and sequence-length statistics for the four training and generation targets. Gap % denotes the percentage of the alignment that is made up of gaps; IDR is the interdecile range between the 10th and 90th percentile of ungapped sequence length.

| Family | MSA depth | Length range | Mean pairwise ID | Gap % | IDR |
|---|---|---|---|---|---|
| GFP (P42212) | 887 | 182–311 | 90.17% | 61.66% | 11.0 |
| NDM-1 (C7C422) | 801 | 186–280 | 43.27% | 57.65% | 26.0 |
| PTP1 (P18031) | 861 | 208–354 | 62.13% | 65.15% | 44.0 |
| FRE (P43127) | 994 | 72–303 | 47.01% | 64.67% | 88.4 |

The four families vary substantially in the mean pairwise sequence identity, with GFPs representing close homologues and FRE containing many more remote sequences. A summary of the properties of sequences selected for training is shown in Table 1.

### 2.3. Model training and initial validation

The model was trained with a maximum of 1000 epochs, with an early stopping patience of 10, measuring validation loss. In practice, this resulted in a total of 20 to 60 training epochs before overtraining commenced. The number of required training epochs also depends on the type of conditioning applied, with conditioning typically increasing the number of epochs needed. The number of model parameters changed with conditioning as well, going from 5 M parameters with no conditioning applied to 5.3 M with both conditioning types applied, length conditioning adding approximately 200k parameters and label conditioning adding 100k.

Training, on average, took under 10 minutes on a single NVIDIA A100 GPU in every configuration, with some variance depending on the family and the type of conditioning applied. Sequence generation throughput averaged 36 sequences per second independent of conditioning, with exact times shown in figure S1. PTP1 has a lower throughput at around 25 sequences per second, due to containing ~ 800 MSA columns, whereas the rest of the families have ~ 600 columns. Conditioning has limited effect on sequence throughput rates, but strongly depends on the size of the input and output. The size of the MSA, on the other hand, has limited impact on the training time.

#### 2.3.1. Effect of sampling concentration

As mentioned above, sequences are generated by numerically integrating the flow of the model toward a target terminal concentration *α*_max_ parameter, which controls how peaked the final per-position amino-acid distribution is (see Methods for further details). We tested *α*_max_ ∈ *{*8, 10, 12*}* and found that increasing *α*_max_ consistently improves structural confidence (structure prediction and validation are described in methods), but at the cost of sequence diversity, as the generated sequences become more similar to the training sequences; see figure S2. For values past *α*_max_ = 12, the model begins to produce duplicate sequences and sequences identical to the training data, see table S1.

The FoldSeek success rate (the fraction of predicted structures matching the expected family fold) rises with increasing *α*_max_, reaching 100% for all four families at *α*_max_ = 12, with the largest increases for GFP and FRE (figure S4). Both NDM-1 and PTP1 have FoldSeek success rates above 90% even for *α*_max_ = 8, whereas GFP and FRE have success rates in the 70s. Increasing *α*_max_ to 10 brings all success rates up to ≥97%, and a final increase to 12 results in a 100% success rate for all four families.

Turning to diversity, for NDM-1 the mean maximum sequence identity to the training set (the identity of each generated sequence to its closest training sequence, averaged over all generated sequences; figure S2) increases from 47% to 60% as *α*_max_ increases from 8 to 12. PTP1 and FRE see similar increases, going from 56% to 71% and 44% to 56%. GFP, which has the highest identity to begin with, goes from 82% to 97%. The pairwise identity distribution among generated GFP sequences also becomes less diverse and more jagged. At *α*_max_ values above 12, the GFP model begins to generate sequences that are both identical to each other and to those in the training data. In summary, increasing *α*_max_ gives sequences that are more confidently predicted to fold correctly, but less diverse, both among themselves and relative to the training set. Unless otherwise specified, the data shown here have been generated with *α*_max_ = 12, as it gave the highest structural success rates across all families.

### 2.4. Sequence diversity and novelty

The sampling-concentration sweep above shows that higher *α*_max_ generates sequences that sit closer to the training distribution. Here we ask whether the model simply reproduces training sequences, or whether generated sequences represent novel points within the sequence space of families. We address this via evaluating the identity to the closest sequence in the training set, and by comparing the overall pairwise identity distribution compared to the training distribution as seen in figure 1.

**Figure 1:**
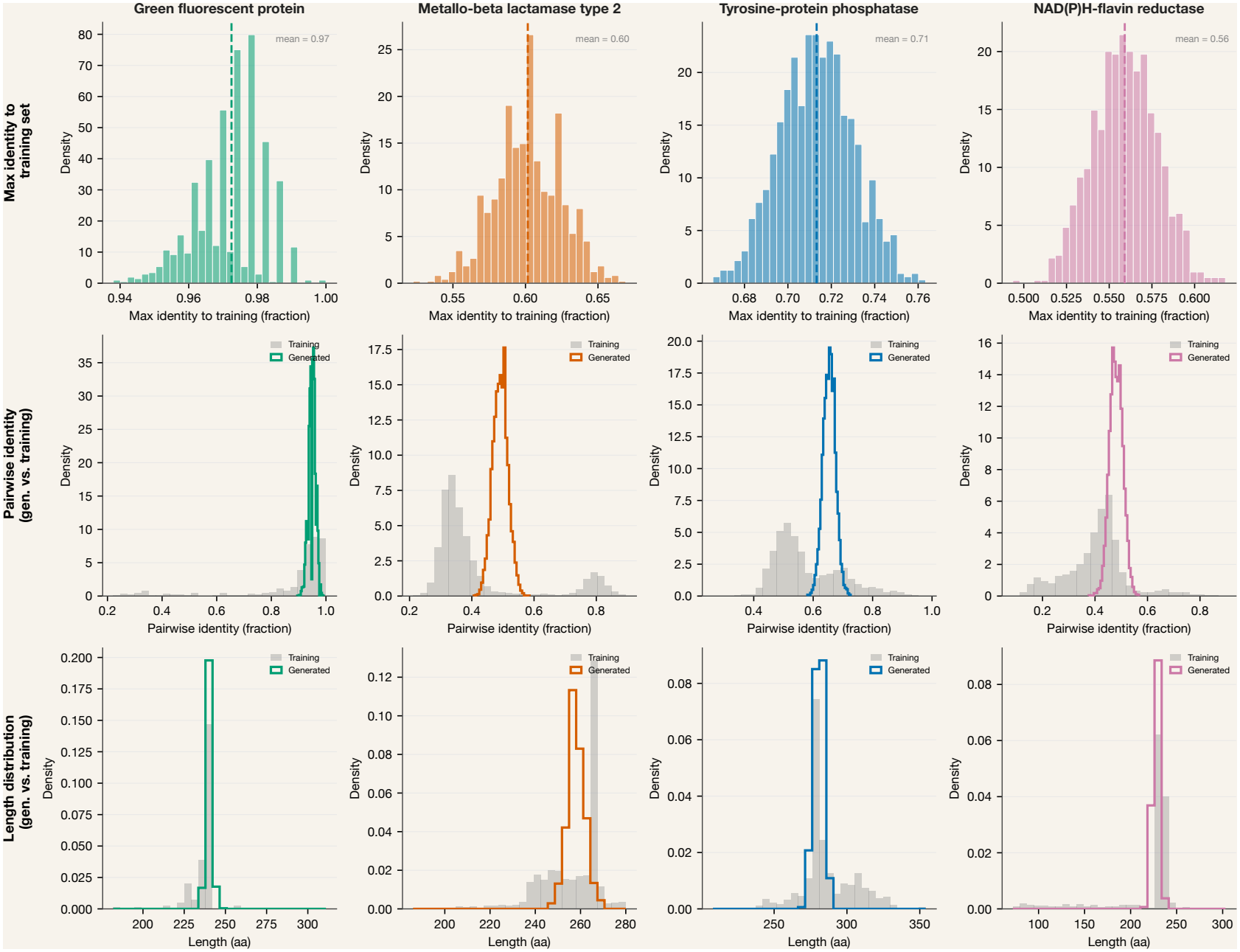
Generated sequences are novel relative to the training set, but match its pairwise identity spread. Top row: distribution of the maximum identity of each generated sequence to any training sequence, for α_max_ = 12 (n = 1000 per family). The dashed line is the mean (GFP 0.97, NDM-1 0.60, PTP1 0.71, FRE 0.56). Middle row: distribution of pairwise identity between all sequence pairs within the training set (grey) and within the generated set (colour). Bottom row: distribution of the generated lengths (colour) overlaid with the lengths within the training set (grey).

GFP, as a more conserved family, has higher training set identity and pairwise identity. The pairwise identity distributions of the generated sequences and the training sequences are almost identical. However, the maximum sequence identity to the training set is quite high, with some sequences nearing 100% sequence identity. This is not unexpected for a highly conserved family and, as discussed in the previous section, can be tuned by modifying *α*_max_.

NDM-1, FRE, and PTP1 are all more diverse than GFP due to being less conserved on the whole. Max identity to training is approximately normally distributed and within the range of the expected pairwise identity of the training data. Looking directly at the pairwise identity, we can see an overall trend that the pairwise identity of the generated sequences is slightly higher than the largest training data cluster. This is especially clear for NDM-1, whose pairwise identity lies between the 30% and 80% cluster seen in the training data. As with GFP, this can be tuned with *α*_max_.

#### 2.4.1. Phylogenetic analysis

While the sequence identity numbers show that we have unique sequences that appear to fit in with the training data distribution, this does not directly tell us how they relate to the training data. To visualise this, figure 2 shows four phylogenetic trees built from a representative subsample of natural family alongside sequences generated with *α*_max_ = 12 (subsampling and tree-construction procedure described in Methods).

**Figure 2:**
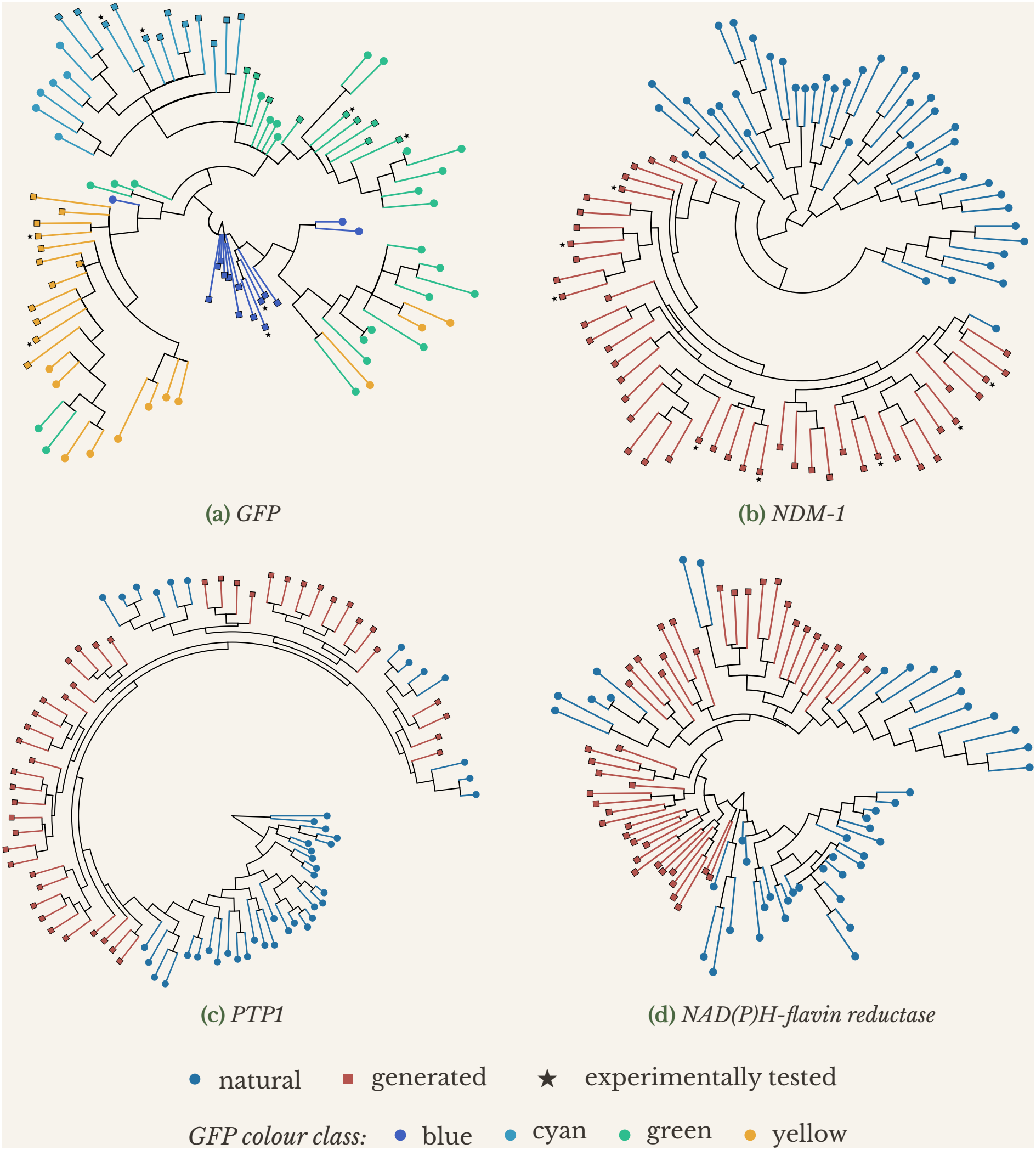
Generated sequences interleave with natural homologues in all four families, but form tighter clusters in NDM-1 and PTP1. Phylogenetic placement of generated sequences relative to natural homologues, for each of the four families (IQ-TREE3, per-family ModelFinder-selected substitution model; see Methods). Circles: natural sequences; squares: generated sequences; stars: experimentally tested candidates. Colours in **(A)** indicate the conditioned spectral class.

**Figure 3:**
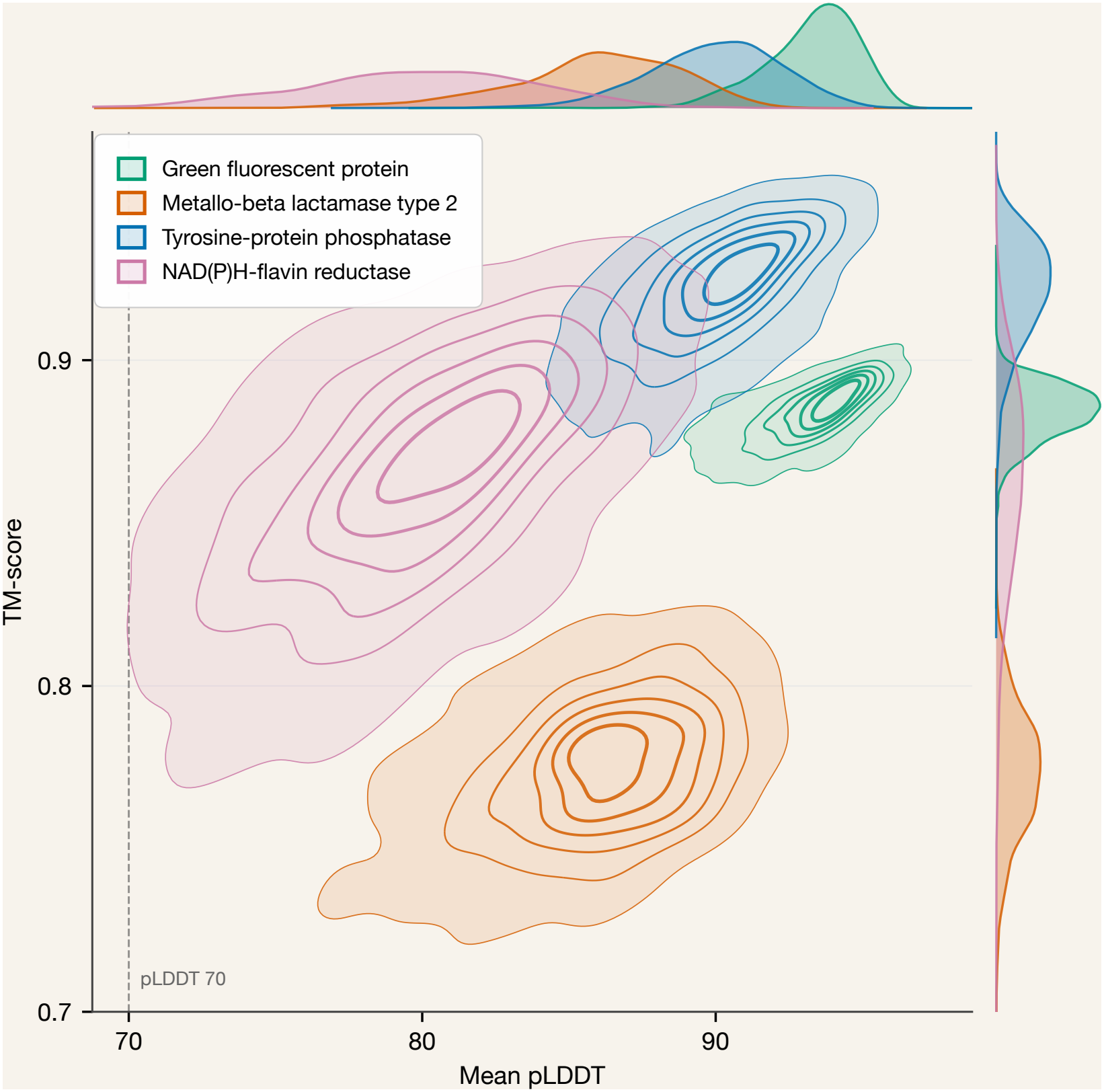
Structure prediction confidence and fold agreement of unconditionally generated sequences (α_max_ = 12). Joint density of per-sequence mean pLDDT (Chai-1) and TM-score to the reference structure, with marginal densities on the top and right (n = 1000 per family). The dashed line marks pLDDT 70. TM-scores are against 1AAX (PTP1, [34]), 1EMA (GFP, [35]), 8GPD (NDM-1, [36]), and AF-P43127-F1 (FRE, [37, 38]). NDM-1 pLDDT excludes the first 40 residues. table 2 reports medians of these distributions.

For clarity, 90 of 887 GFP sequences with <85% sequence identity to wild-type GFP, forming a distinct clade of divergent orthologues, were excluded for legibility. However, all sequences were retained for statistical analysis.

Analysis of the distribution and clustering of sequences in these phylogenetic trees reveals some patterns. Generated FRE and GFP sequences stay close to the natural-level spread; sequences are nearly as diverse as their natural counterparts. PTP1 and especially NDM-1 show a much tighter generated cluster relative to natural sequences. NDM-1 sequences are also more distant from natural sequences than natural sequences are from each other, while being tightly self-clustered, i.e., they are occupying a real but narrow region of family space. This could follow from the bimodal pairwise identity distribution of the NDM-1 training data (peaks at 30% and 80%), while the generated sequences have a single unimodal peak at 50% pairwise identity. PTP1 also has this bimodal training distribution, but less drastic, which may account for the similar behaviour of the two families.

### 2.5. Foldseek results

Next, we were interested in determining whether the sequences generated by the model retained the same fold as the proteins in the desired fold class. To do this, we used FoldSeek, a method that compares a given protein structure with structures in multiple databases and ranks the top matching folds. During validation with a smaller set of 100 generated sequences, predicted by ESM3-open, structures were checked against FoldSeek to ensure correct fold retention[11, 32].

Across the three evaluated *α*_max_ values, FoldSeek domain-match pass rate increased monotonically with *α*_max_ for every family (figure S4A): GFP rose from 72.7% at *α*_max_ = 8 to 100% at *α*_max_ = 12; FRE from 78.8% to 100%; NDM-1 from 94.8% to 100%; PTP1 from 98.9% to 100%. The two families with the weakest *α*_max_ =8 baseline (GFP, FRE) both recovered to *≥*97% by *α*_max_ = 10. This increase tracked structural confidence: pooled across families, at *α*_max_ = 8, sequences with pLDDT below 70 showed a markedly lower pass rate (<60%) than higher-confidence sequences (>90%), whereas this gap narrowed at *α*_max_ = 10 and 12 (figure S4B).

### 2.6. Structural validation across families

While the above FoldSeek provides a high-throughput approach for testing domain-level fold identity with a lightweight structure prediction model, it does not predict whether generated sequences fold with high confidence. Nor does it provide information on whether predicted structures are consistent with experimentally solved structures. We therefore validated a larger set of 1000 sequences per family using Chai-1 with ColabFold-generated MSAs as our main quantitative metric[33]. For ligand-bound complex predictions and selection of sequences for experimental validation, we also used AlphaFold3 as a complementary oracle to enhance confidence in candidate sequences. Structure prediction metrics for the model run without conditioning are shown in table 2. As a reference, we ran 50 natural sequences per family through the same pipeline, 49 sampled at random from the training MSA plus the UniProt reference sequence (see Methods).

**Table 2.** Family pass rates and median TM-score and pLDDT with sample size n = 1000 for α_max_ = 12 for the generated sequences with no conditioning applied, and n = 50 for the natural sequences sampled from the training set. Pass rate is defined as the percentage of sequences with pLDDT > 70 and TM-score > 0.5. NDM-1 has had its first 40 residues excluded from the pLDDT count, as these are frequently truncated for expression due to being unstructured.

| **Family** | **Pass rate** | **pLDDT** | **TM-score** |
|---|---|---|---|
| GFP (generated) | 100.0% | 93.5 | 0.915 |
| GFP (natural) | 100.0% | 95.0 | 0.926 |
| NDM-1 (generated) | 100.0% | 86.2 | 0.803 |
| NDM-1 (natural) | 100.0% | 94.2 | 0.834 |
| PTP1 (generated) | 100.0% | 90.1 | 0.953 |
| PTP1 (natural) | 100.0% | 95.8 | 0.980 |
| FRE (generated) | 99.3% | 80.0 | 0.896 |
| FRE (natural) | 100.0% | 93.4 | 0.964 |

The model generates high-confidence structures that are consistent with the fold of the desired family. All four families pass at *≥* 99.3% with median pLDDT 80.0–93.5 and TM-score 0.803–0.953, as seen in table 2. Natural sequences, run through the same pipeline, score higher (pLDDT 93.4–95.8), with the generated sequences falling short by 1.5–13.4 pLDDT. However, natural sequences are an optimistic reference, as Chai may have seen them in training and may have been able to find deeper MSAs for them.

### 2.7. Conditioning results

The model can additionally condition generation on sequence length and, where labelled training data exists, a discrete functional label. Length conditioning is performed for all four families, and we demonstrate label conditioning using the four GFP chromophore colour classes (green, yellow, cyan, blue), as success can be identified on a sequence level. Furthermore, we evaluate fidelity and success rates at *α*_max_ = 12 across three conditioning regimes: length only, label only, and length + label jointly.

Length conditioning generally achieves reliable control of sequence length for the four families. With *α*_max_ = 12, the mean absolute deviation from the target is 1.9–7.3 amino acids across the four families. However, the performance degrades for out-of-distribution lengths and lengths with sparse training support, meaning fewer than 5% of sequences are within *≤* 10 amino acids of the target length; see figure 4. The deviation is larger for targets with sparse support (4.6–26.0 amino acids) and largest for targets entirely outside the training length range (14.9–86.4 amino acids). The performance for other values of *α*_max_ can be seen in figure S3.

**Figure 4:**
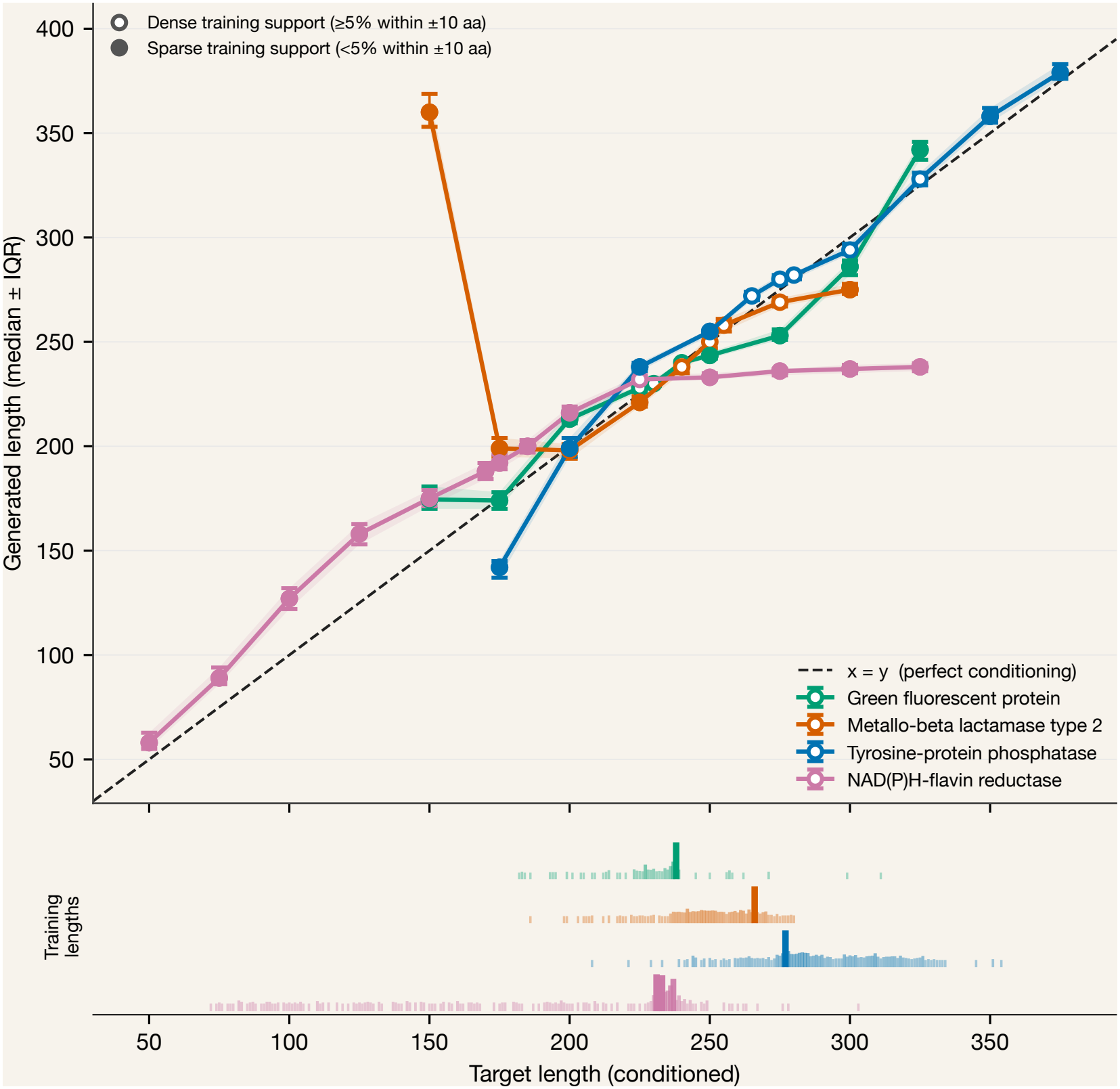
Length conditioning is accurate in length ranges where there exists supporting training data. Median generated length ( ±IQR) against the requested length at α_max_ = 12 (n = 150 sequences per target length). The dashed line is perfect conditioning. Open markers: dense training support (≥ 5% of training sequences within ±10 residues of the target length). Filled markers: sparse training support, or no training support. The rug below shows each family’s training length distribution; tick height and opacity scale with sequence count.

In comparison, unconditional generation does not fully reproduce the training-set length distribution, with generated sequences shifted to longer lengths and with greater spread across all four families. However, it gets close with high *α*_max_, see figure S2. Length conditioning thus allow sampling of less frequently observed sequence lengths that are rarely sampled in unconditional sampling.

Label conditioning reaches high accuracy (98–99%) for every label, independent of how often the label occurs in the training data (2.4% for blue, 66.2% for green; figure 5). Sequence quality is also similar across labels, with a mean pLDDT of 93–94.

**Figure 5:**
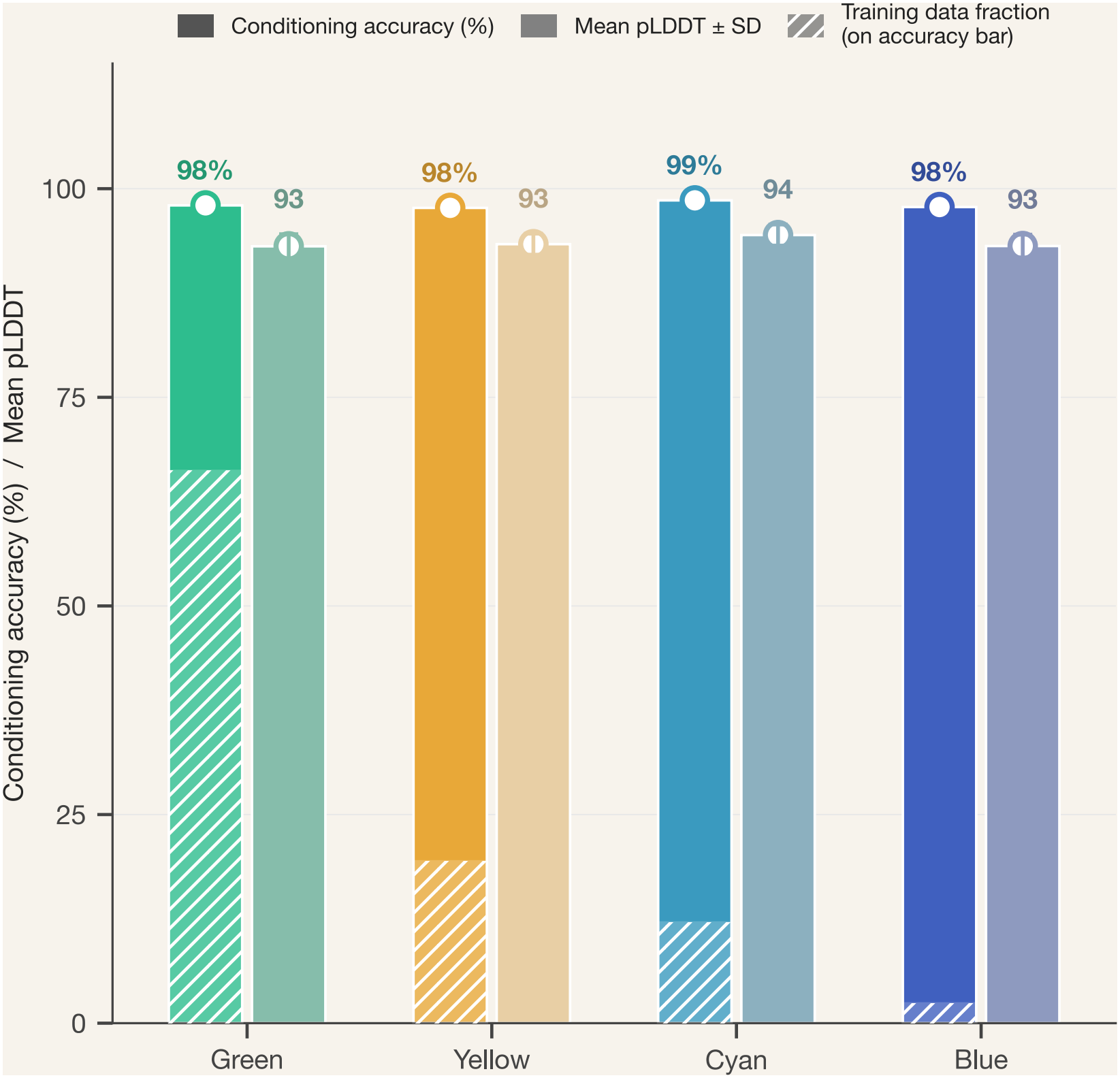
Label conditioning reaches 98–99% in silico accuracy for every GFP colour class, including the rarest. Solid bars: percentage of generated sequences whose colour class, assigned from chromophore-region residues (table S3), matches the requested label. Hatches part: fraction of the training MSA with the given label (green 66.2%, blue 2.4%). Light bars: mean pLDDT± SD from Chai-1. Conditioning was label only, no length target; α_max_ = 12; n = 1000 per label. Accuracy is a sequence-based prediction, not based on experimental data.

**Figure 6:**
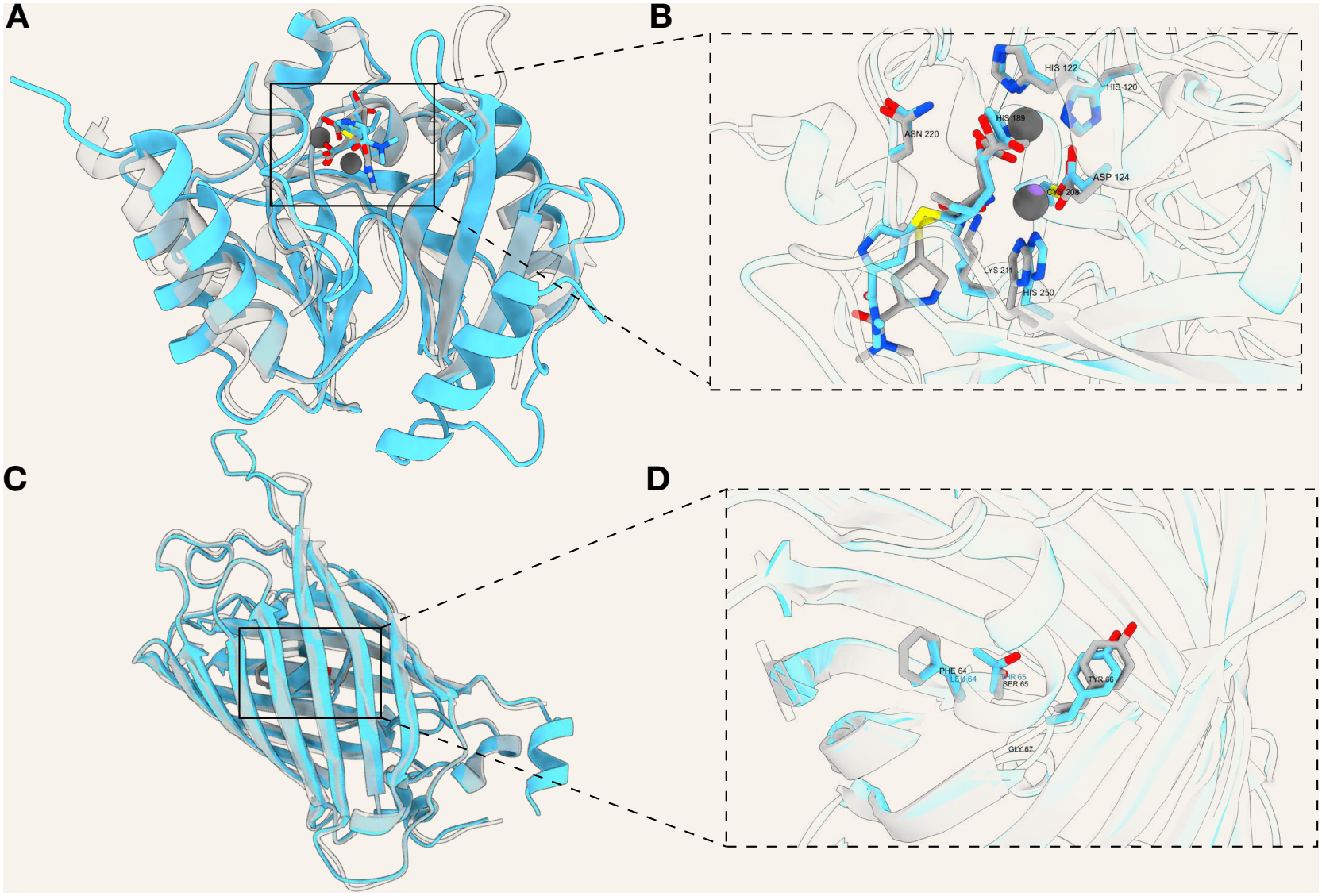
Overlay of top Chai-1 predicted candidate structures of NDM-1 and GFP (blue), overlaid with WT (grey). **A-B)** Structural overlay of the best predicted NDM-1 candidate complex (blue) with the crystal structure 4EYL (grey)[39]. **B)** shows the active site comparison, where all residues correspond to each other. The fiexible tail of the ligand meropenem differs between the WT and the prediction. **C-D)** Structural overlay of the best predicted GFP candidate with the WT crystal structure 1GFL[20]. D shows the chromophore residues overlaid, though the designed variant is an eGFP variant and contains the mutations F64L and S65T.

**Figure 7:**
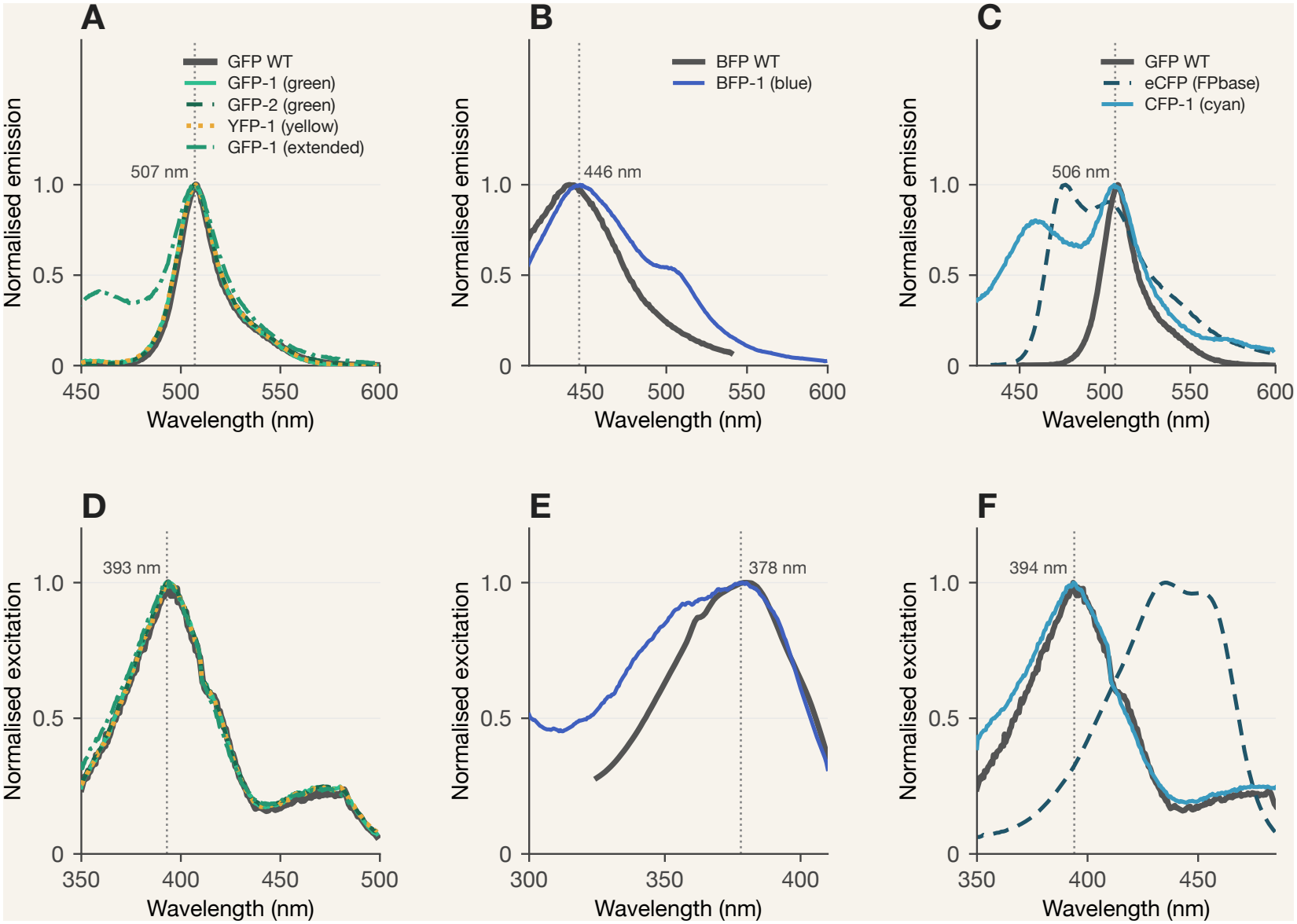
Normalized fluorescence emission and excitation spectra of GFP, YFP, CFP and BFP designs overlaid with their respective wild-type (WT). (A–C) Normalised fluorescence emission spectra of the GFP designs (A), BFP design (B) and CFP design (C), overlaid with their respective WT reference spectra. The emission spectrum of BFP-WT and eCFP-WT was obtained from FPbase [40]. (D–F) Normalised fluorescence excitation spectra of the GFP (D), BFP (E) and CFP (F) designs. The wavelengths corresponding to the main emission and excitation maxima are indicated in the respective panels.

Co-conditioning on GFP colour and length keeps the length control of length-only conditioning, but label accuracy depends on whether the requested combination occurs in the training data (figure S5). The mean absolute deviation from the target length is 1.7–2.6 amino acids for targets with dense training support and 10.3–14.9 amino acids for targets outside the training range, compared with 1.9 and 15.4 amino acids for length-only conditioning. Label-accuracy is near 100% for all four labels at targets of 230–250 amino acids, where all labels have training examples. Outside this range, it falls for labels that are rare at those lengths, most clearly for blue, which drops to 0% from a target of 275 onward while still producing sequences within about 15 amino acids of the requested length, whereas green stays above 95% up to a target of 325.

### 2.8. Candidate selection and ligand-binding validation

To identify individual designs for experimental follow-up, we selected candidates by predicting generated sequences with each family’s physiologically relevant ligands and ranking by a composite score combining structural confidence with correct ligand-contact geometry (exact procedure described in Methods). We examine this process in detail for NDM-1, where a ligand bound crystal structure allows direct comparison against a known ligand-binding pose.

Two families were selected for experimental characterisation, GFP and NDM-1. GFP was chosen because its activity and fluorescence depend on the internal geometry of the fluorophore and beta barrel, which made it suitable for testing whether Syzygy recapitulates functional internal geometry in proteins. NDM-1 was chosen as its function requires coordination of two Zn ions and a substrate, meaning the binding pocket does not simply need to be conserved, but the overall geometry must lead to an active enzyme. These two families allow for measurement of both intrinsic and extrinsic properties of the proteins.

Selection of the NDM-1 candidates involved prediction with two Zn ions and a meropenem substrate. The meropenem substrate, however, has a flexible tail, which complicated prediction. Both Chai-1 and AlphaFold3 fail to predict the pose found in the crystal structure when predicting the native complex: the Zn ions and the front of the meropenem are in the correct position, but the pose of the tail differs. This tail region is where one of the binding pocket residues (K211) makes contact with the substrate. This residue contact was most frequently missed in the predictions of the generated sequences; however, full binding pocket contacts remained a hard requirement for any experimental candidate.

While selection for candidates for NDM-1 allowed sequences from all tested *α*_max_ values, all of the top candidates ended up selected from *α*_max_ = 12. Looking at Chai-1, 76% of predictions passed the requirement to contain contacts for all binding pocket residues, with all failures missing the same residue: K211. Contacts for the Zn ions and the “head” of the meropenem were fully retained in every prediction. The mean pLDDT for the predictions was 82.6, with the protein and Zn ions being the most confidently predicted, and meropenem lowering the average with pLDDTs in the 70s. The mean TM-score of the protein predictions themselves when compared to the crystal structure was 0.852, indicating the correct fold. AF3 mostly recapitulated these results and agreed with Chai-1 for passing candidates, with some minor numerical differences (72% having full contact, mean pLDDT 86.9, mean TM-score 0.843).

Unlike NDM-1, where all experimental candidates were drawn from *α*_max_ = 12, for the GFP candidate set we deliberately included sequences from *α*_max_ = 10 (6 out of 8 candidates) alongside *α*_max_ = 12 to prioritise sequence diversity. No sequence with sequence identity *≥* 95% to the training set was chosen for experimental validation.

### 2.9. Experimental validation

#### 2.9.1. Fluorescence measurements

The GFP designs were recombinantly expressed in *E. coli* and purified with metal affinity chromatography via their His-tags (details in Methods). Overall, the recombinant proteins showed lower expression levels than the native GFP used as the wild-type reference. Nevertheless, we confirmed the proteins in the elution fraction by Western blotting S6. To functionally characterize the artifical homologs we characterized fluorescence emission and excitation spectra. YFP-2 and CFP-2 did not show detectable fluorescence under the conditions tested. This was consistent with the low amount of protein detected in their respective elution by Western blot analysis. The GFP designs, as well as YFP-1, displayed excitation and emission spectra characteristic of WT GFP. WT GFP showed an emission maximum at 507 nm, which was identical to that observed for GFP-1 and YFP-1. GFP-2 showed a slightly shifted emission maximum at 506 nm. Similarly, the excitation maximum of WT GFP was observed at 393 nm, while the designs showed maxima within the range of 392–394 nm. Overall, the excitation and emission maxima of the designs were comparable to those of WT GFP. However, YFP-1 did not display distinct excitation or emission peaks in the YFP spectral range.

CFP-1 showed its main excitation and emission maxima within the GFP spectral range, at 394 nm and 506 nm, respectively. In addition, a second emission peak was observed at 460 nm, which was shifted towards shorter wavelengths compared with the typical CFP emission peak at approximately 480 nm.

Experimental studies have shown that the flexible C-terminus have the ability to improve the folding of GFPs[41]. To test whether that is the case in our system, we tested a version of GFP-1 with a restored C-terminal (extended GFP-1) region that had previously been truncated. This did, however, not show an increase in expression. On the other hand, the extended GFP-1 displayed a spectral profile that differed from the other GFP constructs, showing an additional emission peak at 460 nm, similar to CFP-1. The excitation spectra of extended GFP-1 showed an excitation peak at 395 nm, corresponding to the excitation maximum for GFP WT.

The BFP construct showed an emission peak at 446 nm, consistent with fluorescence in the blue region. In addition, a second emission peak was observed at approximately 508 nm, corresponding to the emission region of GFP. This second peak was not observed in the BFP WT control obtained from FPBase[40, 42]. Detailed emission and excitation peaks can be seen in the supplementary S2.

#### 2.9.2. NDM-1 activity

Recombinant expression of NDM-1 did not result in any substantial amount of purified protein. Since natural NDM-1’s are typically expressed with a solubility tag we constructed genetic constructs with an N-terminal SUMO tag [17, 25]. Adding the SUMO tag enabled protein expression, however expression levels remained lower than those of native NDM-1. To investigate whether redesigning surface residues could improve the expression of the generated proteins, we redesigned the solvent-exposed residues of NDM-D2, our NDM-1 design with the best *in silico* metrics. This redesigned variant was also fused to the N-terminal SUMO tag. However, the surface redesign did not result in an improvement in protein expression.

Purification did not result in pure protein, it revealed co-purification of molecular chaperones, indicating difficulties with folding of the protein in the cells. The presence of the target proteins in the purified fractions was confirmed by Western blot analysis S6. Interestingly, two distinct bands were observed for NDM-D5, NDM-D6 and NDM-D7, suggesting a partial cleavage of the protein. The upper band, at approximately 40 kDa, corresponded to the expected molecular weight of the His-SUMO-tagged constructs, whereas a second band at approximately 35 kDa was also detected.

Due to the low yield and the presence of co-purified proteins, enzyme activity measurements were challenging. Nevertheless, detectable activity was observed for NDM-D6 and NDM-D7. Both variants showed approximately one-third of the WT activity after normalization to protein abundance figure 8.

**Figure 8:**
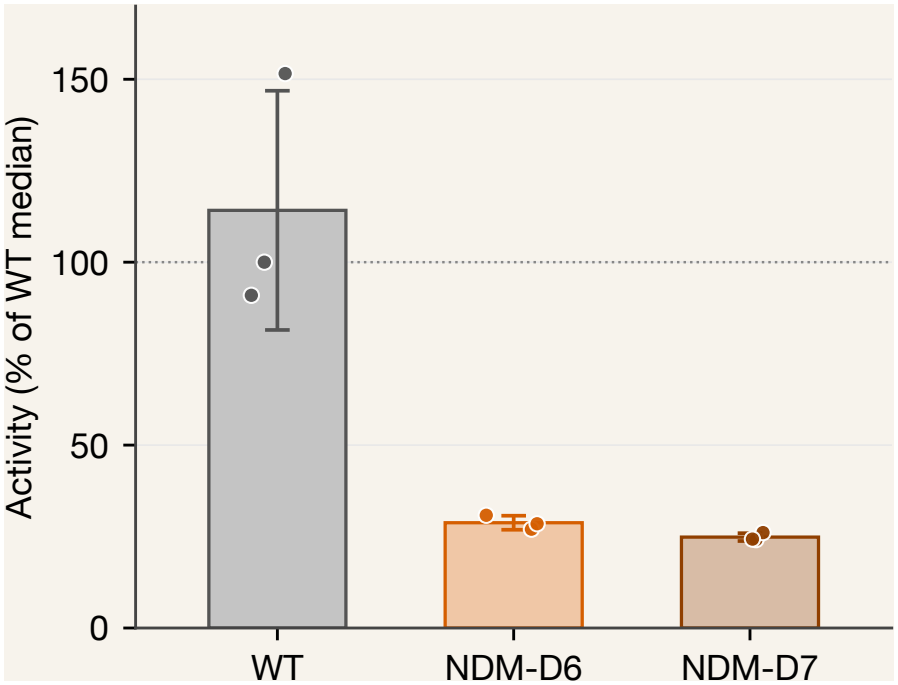
Activity of NDM1 Designs normalized by concentration and WT activity. The activities of NDM-D6 and NDM-D7 were normalized to protein concentrations determined by Western blot analysis. The normalized activities were subsequently expressed relative to WT activity, with the WT median set to 100%.

## 3. Discussion

Here, we present Syzygy, a generative model for creating artificial homologues of protein families as starting points for engineering new protein functions. Many protein folds support a broad range of biological activities, making them attractive frameworks for functional diversification. Through evolution, protein families have accumulated sequence features that encode not only foldability and stability, but also functional properties such as conformational dynamics, substrate-binding pockets, and allosteric regulation. By learning these family-level sequence patterns, generative models can explore diverse regions of sequence space while preserving the underlying features that support protein structure and function. The resulting artificial homologues can provide versatile starting points for engineering proteins with altered substrate specificity, catalytic activity, stability, expression, and molecular interactions. The method is also well suited to generate realistic and diverse sequences for the creation of synthetic MSAs and phylogenetic trees, with many applications in evolutionary research.

The lightweight nature of the generative model enables rapid training on individual protein families, allowing models to be tailored to specific research questions and design objectives. Furthermore, the model can leverage homologous sequences that might otherwise be excluded from training datasets. Large protein language models, such as ESM-2, often employ sequence clustering and subsampling to reduce redundancy during training[43]. In contrast, our Syzygy can be trained on deep alignments containing highly similar sequences, allowing it to exploit the sequence information available within individual protein families.

Beyond its flexibility in training, the model also enables control over the diversity of generated sequences through a single parameter, *α*_max_. Lower values promote exploration of more distant regions of sequence space, generating sequences that are more divergent from natural homologues and from one another. This increased diversity is accompanied by a reduction in average predicted structural confidence, although highly confident structures can still emerge from the more diverse sequence pools. The choice of *α*_max_ therefore provides a means of balancing sequence exploration against structural confidence, depending on the protein family and the intended design objective. For families with limited natural sequence diversity, such as GFP, lower *α*_max_ values can promote greater sequence exploration. In contrast, for more diverse families such as NDM-1, *α*_max_ can be selected to favour sequences with the desired binding-pocket geometry.

Protein families often exhibit substantial variation in sequence length, particularly in terminal regions and loops, which can influence substrate accessibility, specificity, conformational dynamics, and protein expression[44]. Exploring length variation therefore offers an additional dimension for engineering protein properties and functions. However, many generative models require a fixed sequence length during inference, limiting their ability to explore this dimension of sequence space. In contrast, Syzygy supports variable-length sequence generation, both through unconditional sampling and explicit conditioning on sequence length. Unconditional generation produces an approximately normal distribution of sequence lengths, with limited sampling of the extremes of the natural length distribution. By explicitly conditioning on length, the model can target these less frequently sampled regions, with a strong correlation between specified and generated sequence lengths. However, the model is limited to interpolation within the length range represented in the training data and cannot reliably generate sequences beyond this range.

Conditional sampling provides a means to generate sequences that consistent with a subgroup within an alignment, while still utilizing the overall patterns found in the family. The specific function of a protein within a family is often encoded at a few positions, so conditioning on a functional label concentrates changes at these positions while generating sequence features consistent with the family’s overall sequence patterns. There are few BFP sequences in the training sequences, and they are not sufficient to separately train the model on. Nonetheless, a model conditioned on blue chromophores almost exclusively samples BFP-like sequences.

The multiple sequence alignment (MSA) is both a key determinant of model performance and an important means of controlling the properties of generated sequences. Consequently, alignment curation requires consideration not only of the number and diversity of sequences, but also of their biological characteristics and relevance to the design objective. For NDM-1, model performance improved with larger alignments, despite a reduction in overall sequence identity. In contrast, GFP generation quality improved when sequences derived from red fluorescent proteins (RFPs) were excluded, despite their shared evolutionary ancestry. These observations highlight the importance of family-specific MSA curation and suggest that selecting sequences based on biological properties, such as organism of origin or known functional activity, could provide an additional means of tailoring generated sequences toward specific engineering objectives.

The experimental results demonstrate that Syzygy can generate functional artificial homologues that provide starting scaffolds for further optimisation and functional diversification. However, the generated designs appear to inherit not only the functional characteristics of their parent protein families, but also some of their limitations, including difficulties with expression. For example, wild-type NDM-1 is difficult to express, and truncation of its N-terminal 38 residues has previously been shown to increase expression yields approximately 50-fold [27]. Since much of the length variation in our NDM-1 designs occurred in the N-terminal region, differences in this region may have contributed to their reduced expression. Although we truncated the designs to approximate previously successful N-terminal constructs, differences in their N-terminal sequences and lengths may require design-specific truncation strategies. Similarly, the GFP designs contained relatively extensive loop insertions, which may have contributed to their expression difficulties. These observations highlight the importance of considering family-specific characteristics when curating training MSAs and selecting designs for experimental validation.

Despite these expression challenges, several designs retained biological function. The GFP designs largely exhibited fluorescence emission characteristics similar to wild-type GFP, indicating successful chromophore formation despite reduced expression [45]. The NDM-1 designs showed more pronounced expression difficulties and required an N-terminal SUMO tag to enable detectable expression. Co-purification of molecular chaperones suggested potential folding or stability limitations, while Western blot analysis revealed additional lower-molecular-weight bands for NDM-D5, NDM-D6, and NDM-D7, potentially indicating proteolytic cleavage. Previous studies have reported C-terminal truncation of NDM-1 under destabilising conditions, including zinc depletion [46], which could explain the observed band patterns. Nevertheless, NDM-D6 and NDM-D7 retained approximately one-third of wild-type enzymatic activity, demonstrating that the generated sequences can preserve catalytic function despite substantial sequence divergence and expression difficulties. These results also highlight the limitations of computational structure predictions, which provide measures of fold plausibility rather than experimental functionality. Although pLDDT and TM-score can inform the selection of promising designs, they do not reliably predict expression, stability, or activity. Overall, Syzygy provides a rapid means of generating functional protein scaffolds that can serve as starting points for subsequent optimisation through experimental screening and iterative protein engineering.

The model can be investigated further, both in how it is used and in what it can do. Length conditioning can be used to build loop libraries: sequences with a range of target lengths can be assayed to relate loop length to activity and expression and to see how different loop lengths affect certain families. Setting *α*_max_ to a high value can be used for generating libraries, as many positions will be effectively fixed and the generated sequences will be very close to natural. Because *α*_max_ can be changed between training rounds, it can also serve as the exploration parameter in an active learning loop, with the assay results of each round informing the next library. Label conditioning has so far been demonstrated on GFP colour. Applying it to other families and to other kinds of labels, such as thermostability, host organism and sub-family, would show which properties the model can be steered towards. Continuous properties, such as melting temperature, could be binned into classes. Sampling could also be biased towards or away from chosen amino acids, for example, to limit cysteines, which were removed here by filtering after generation. Classifier-free guidance could be implemented to strengthen conditioning, particularly for lengths near the ends of the training range. A first attempt at this did not improve performance, and it may need its own hyperparameter tuning for optimal performance. Finally, ESM3 or ESMC embeddings could be added as a conditioning input, which may improve sampling for small families or allow smaller MSAs, in exchange for a pretrained-model dependency and a larger computational footprint.

## 4. Data Availability

The source code for Syzygy is available at https://github.com/Andre-lab/Syzygy.

## 5. Acknowledgement

This work was supported by the Wallenberg AI, Autonomous Systems and Software Program (WASP) funded by the Knut and Alice Wallenberg Foundation to IA. This work was funded by a grant to IA from the Swedish Research Council, project number 2022-04221.

## 6. Materials and Methods

### 6.1. Dirichlet Flow Matching for Protein Sequences

Dirichlet Flow Matching extends continuous normalising flows to data residing on a probability simplex Δ^*K–*1^, where *K* represents the alphabet size, which is 21 in this case[13]. Protein sequences from an aligned MSA were represented as sequences of tokenised vectors *x* ∈ *{*0, 1, …, 20*}*^*L*^, where each amino acid from the 21-character alphabet (20 standard amino acids plus gap “-”) is mapped to a unique integer index, and L is the sequence length including gaps. These discrete states are treated as vertices of the (*K –* 1)-simplex.

The training objective defines a probability path *p*_*t*_(*x*) that interpolates between a Dirichlet prior distribution at t=0 and the data distribution at t=1. Following the original paper[13], we trained the neural network in Syzygy to approximate the vector field *v*_*θ*_(*x*_*t*_, *t*) driving this probability path. The network was optimised using the cross-entropy objective:

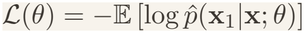

where *θ* denotes the network parameters, *x*_1_ the final data being predicted, and 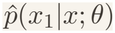 the model’s predicted probability over amino acids at each position given the noisy state *x*. It can be shown that this has the same minimiser as the traditional *L*_2_-like conditional flow matching loss, and is thus a valid objective, and allows for the vector field to not have to be calculated during training[13]. Before being passed to the network, the noisy simplex sample *x*_*t*_ *∈* ℝ^*K*^ is expanded to 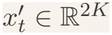 by concatenating it with a copy of itself reweighted by a prior pseudocount, as per the original implementation.

### 6.2. Model Architecture

The vector field *v*_*θ*_ was parameterised using a Transformer network with AdaLN-Zero conditioning[14]. The model was implemented in PyTorch, and its parameters can be seen in table 3. Instead of learning static scale and shift parameters (as in standard LayerNorm), this model regresses them from the conditioning information. The expanded input 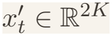 is linearly projected to the hidden dimension *D* and combined with a fixed sinusoidal positional encoding, giving the sequence tensor *x∈* ℝ^*B×L×D*^ used throughout the network. The architecture choices can be seen in figure 9.

**Figure 9:**
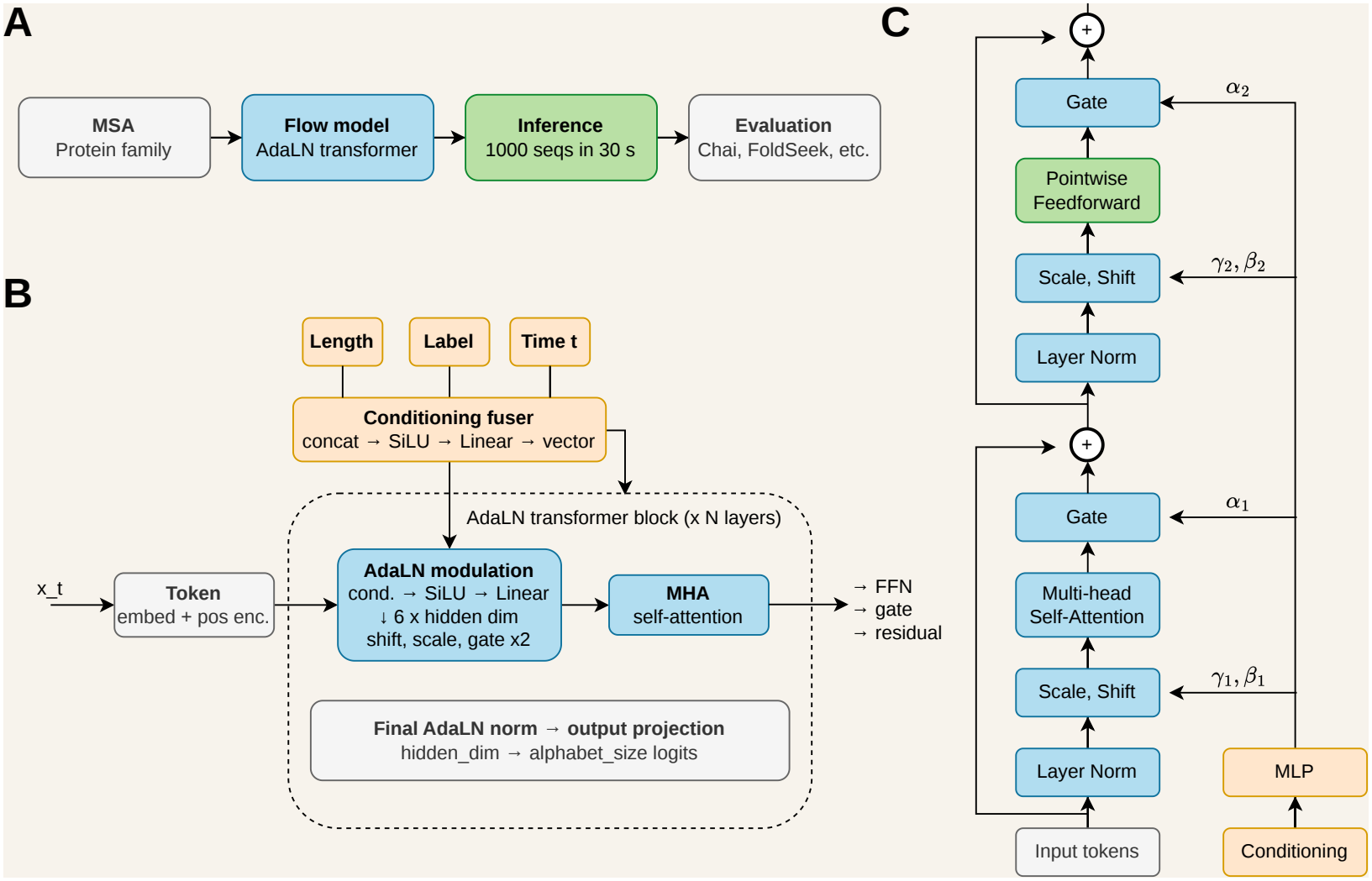
Overview of the “ow-matching architecture. **(A)** Training and inference pipeline: an MSA for a given protein family is used to train the Syzygy AdaLN transformer flow model, which is then sampled (1000 sequences in 30 s) and evaluated with downstream structure/quality tools (Chai, FoldSeek, etc.). **(B)** The AdaLN transformer model. Length, label, and time conditioning signals are pre-embedded before being fused into a single conditioning vector, which modulates each of the N stacked AdaLN transformer blocks alongside the embedded, position-encoded input x_t_. A final AdaLN norm and output projection map the last hidden state to per-position logits over the alphabet. **(C)** Detail of a single AdaLN transformer block, showing the scale/shift modulation (γ, β) before each of the self-attention and feedforward sub-layers, and the gate (α) applied to each sub-layer’s output before the residual connection.

**Table 3.** Model hyperparameters and their values.

| **Parameter** | **Value** |
|---|---|
| Dropout | 0.2 |
| Layers | 4 |
| Heads | 8 |
| Hidden dimension | 256 |
| Learning rate | 0.0005 |
| Batch size | 128 |

**Table 4.** Emission and excitation spectra for fluorescent proteins.

| Fluorophore | Emission range | Emission peak | Excitation range | Excitation peak | Slit width |
|---|---|---|---|---|---|
| GFP | 450 - 600 | 508 | 350 - 500 | 395 | 5 or 15 |
| BFP | 400 - 600 | 445 | 300 - 400 | 380 | 15 |
| CFP | 450 - 550 | 435 | 300 - 400 | 447 | 5 |
| YFP | 500 - 600 | 527 | 400 - 500 | 514 | 5 |

- *x∈*ℝ^*B×L×D*^ is the resulting embedded, position-encoded sequence tensor
- *c∈*ℝ^*B×D*^ is the conditioning vector derived from time *t* and, depending on the conditioning regime, sequence length and/or a discrete label

Here, *B* is the batch size, *L* is the sequence/MSA length, and *D* is the hidden dimension.

The architecture supports two independent conditioning signals in addition to time: sequence length and a discrete label, each toggled independently at model construction. Label conditioning was trained for the GFP family only, using the fluorescence colour class as the label, under all four combinations of the two signals: unconditioned, length-conditioned, label-conditioned, and label+length-conditioned. The remaining three families were trained unconditioned and length-conditioned only.

Each active conditioning signal is first embedded independently. Time *t* is mapped to *D* dimensions using random Gaussian Fourier features, followed by a small MLP. Sequence length is normalised to [0, 1] using the minimum and maximum ungapped lengths seen in training, passed through a tanh squashing function, then projected to *D* dimensions by a two-layer MLP. The discrete label, when used, is mapped to *D* dimensions via a learned embedding lookup. When more than one signal is active, the individual embeddings are fused by a small MLP into a single conditioning vector before being passed to AdaLN.

The conditioning vector *c* is passed through an MLP to predict six modulation parameters per dimension:

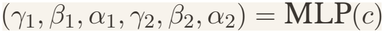

where *γ* represents the scale, *β* represents the shift/bias, and *α* represents the gate (scaling factor for the residual connection). Indices 1 and 2 refer to the attention and feed-forward blocks, respectively.

Standard LayerNorm normalises the input *x* to have zero mean and unit variance. In AdaLN, we modulate this normalised vector using the predicted scale and shift. The code uses (1 + *γ*) to ensure that the MLP does not output zeros if *γ* is zero, which it may be as it is initialised to zero.

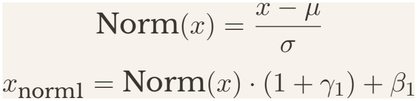

Then, the modulated input enters the multi-head self-attention (MHSA) block. The output is scaled by gate *α*_1_ before being added back into the original input *x* (residual connection).

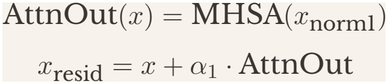

This process then repeats for the Feed-Forward Network (FFN).

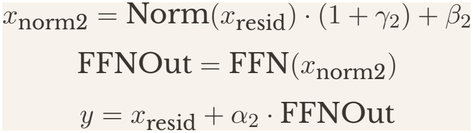

where *y* is the output of the block.

After the final block, a separate AdaLN step predicts only a scale and shift (*γ*_*f*_, *β*_*f*_)= MLP(*c*), without a gate since there is no residual connection to the output. This is applied to Norm(*y*) in the same way as *x*_norm1_. A final linear layer then projects the result from *D* to the alphabet size *K*, giving the output logits.

### 6.3. Training Procedure

The model was trained on four families: the hydrolase family Tyrosine-protein phos-phatase non-receptor type 1 (P18031, PTP1), the oxidoreductase family NAD(P)H-flavin reductase (P43127, FRE), the hydrolase family Metallo-beta-lactamase type 2 (C7C422, NDM-1), and the green fluorescent protein family (P42212, GFP). The model was trained with the Adam optimiser for 1000 epochs, or until validation loss had not decreased for 10 epochs.

### 6.4. Sequence Generation

Sequences were generated by sampling *x*_0_ from the Dirichlet prior and solving the probability flow ODE with an Euler solver over 100 integration steps. Following Dirichlet Flow Matching, the path is parameterised by the Dirichlet sampling concentration parameter *α*: integration proceeds from *α*_min_ = 1, the prior with no sequence information, to *α*_max_. *α*_max_ sets the endpoint concentration of the sampling path and therefore how peaked the terminal per-position amino-acid distribution is. A higher *α*_max_ gives a more peaked terminal amino-acid distribution at each position. Three values were evaluated for each trained conditioning regime: *α*_max_ ∈ *{*8, 10, 12*}*, where 8 is the default used by the original Dirichlet flow matching paper[13]. Values above 12 were not pursued further after initial testing showed generated GFP sequences increasingly replicating training-set sequences.

### 6.5 Structure Prediction and Validation

Generated sequences were initially predicted with ESM3-open for quick validation[11]. Structures were then checked against FoldSeek to ensure the correct fold[32]. Final predictions were performed using Chai-1, both with and without MSAs, using family-specific ligands for binding validation[33].

#### 6.5.1. FoldSeek Structural Search and Classiflcation

For each of the *n* = 100 generated sequences evaluated per family and *α*_max_ condition, the predicted structure (predicted by ESM3-open [11]) was submitted to the FoldSeek search server and queried against nine reference databases: AlphaFold/UniProt50, AlphaFold/Swiss-Prot, AlphaFold/Proteome, PDB100, CATH50, the ESM Metagenomic Atlas (mgnify_esm30), the Big Fantastic Virus Database, BFMD, and GMGCL[32]. Matches with alignment probability *≥* 0.5, E-value *≤* 10^*–*4^, and gap fraction *≤* 30% were retained as high-confidence hits. No pre-filtering by predicted structural confidence (pLDDT or TM-score) was applied before FoldSeek submission for the reported *α*_max_ comparisons, so every generated sequence in the evaluated set received a FoldSeek search regardless of its structure-prediction confidence.

For each high-confidence match, the target’s Pfam and InterPro domain annotations were resolved via the UniProt and PDBe REST APIs[47], subject to a fixed per-sequence lookup budget to keep query time tractable. The most abundant matches were resolved first; because family-conserved folds return overwhelmingly correct-family hits among their top matches, this budget affects the precision of reported domain-frequency percentages but not the pass/fail call for individual sequences. A sequence was scored as passing the FoldSeek structural validation if at least one classified high-confidence hit shared a Pfam or InterPro identifier with the sequence’s target family.

#### 6.5.2. Prediction of Natural Sequences

For each of the four families (GFP, NDM-1, FRE, PTP1), gaps were removed from the family’s training MSA, the sequences were shuffled (seed 1), exact duplicates were removed, and 49 sequences were kept. The UniProt reference sequence for the family (P42212, C7C422, P43127, P18031) was added to each sequence subset, giving 50 sequences per family. No filtering by sequence identity, length, or taxonomy was applied. Structures were predicted with Chai-1 using MSAs from the ColabFold MSA server and ESM embeddings, with default sampling settings (3 trunk recycles, 200 diffusion timesteps, 5 samples per sequence) and no ligands; the same procedure as for the generated sequences.

### 6.6. Multiple sequence Alignment

Multiple sequence alignments (MSAs) were generated separately for each protein family. A reference protein sequence was selected from UniProt and used to find homologous sequences from protein BLAST (Blastp). Blastp searches were performed using the non-redundant protein sequences (nr) database with a maximum of 5000 sequences. The resulting sequences were filtered before alignment. Substantially truncated sequences were removed to increase the quality of the alignment. To reduce sequence redundancy, sequences from three of the protein families were clustered using CD-HIT with a sequence identity threshold of 90% (-c 0.9), and one representative sequence was retained from each cluster. For the GFP dataset, a sequence-identity filtering of 98% was applied, as applying the 90% identity threshold substantially reduced the number of available sequences due to the high sequence similarity within this family. The remaining sequences were aligned using MAFFT.

### 6.7. Phylogenetic Analysis

For each protein family, training sequences and generated sequences were subsampled and combined into a single reduced sequence set for tree construction. Training sequences were reduced from the full training MSA in three stages:

1. GFP sequences with less than 85% sequence identity to wtGFP were excluded, as these sequences tended to be from different organisms and formed a large separate clade which made tree legibility worse. This resulted in 90 sequences out of 887 being removed.
2. Near-identical sequences were removed using CD-HIT at 98% identity [48].
3. Sequences were sampled using farthest-point sampling on ESM2-embeddings (35M parameter version) of the sequences to select 45 diversity-representative sequences per family[43]. Farthest-point sampling was performed starting at a random sequences and then sampling the rest using Euclidian distance, greedy selection, and a fixed seed.

The generated sequence set for tree construction combined an experimental candidate set for GFP and NDM-1, with supplements drawn from the broader *α*_max_ = 12 pool ranked by pLDDT, balanced across colour-conditioning classes for GFP, for a total of 45 generated sequences per family. For PTP1 and FRE, the entire generated set was pulled from their respective pLDDT-ranked *α*_max_ = 12 sequence pool.

Generated sequences were threaded onto the frozen natural MSA using MAFFT( `mafft --add --keeplength`), preserving the natural sequences’ original alignment columns[49]. Maximum-likelihood phylogenetic trees were inferred with IQ-TREE3 (`iqtree3 -s COMBALIGNMENT.fasta -m MFP -B 1000 -alrt 1000 -bnni -T AUTO -ntmax 8 -seed 12345`), with the substitution model selected independently per family using ModelFinder[50]. Trees were visualised with ETE3 as midpoint-rooted, for display purposes only[51].

To quantify sequence divergence beyond what the tree topology shows visually, pairwise patristic (tree-path) distances were computed between all leaf pairs on the untransformed tree and grouped by sequence provenance (natural-natural, natural-generated, generated-generated). Distributions were compared using two-sided Mann-Whitney U tests.

### 6.8 Ligand Binding Prediction

All sequences were predicted with ligands using Chai-1 with MSAs, as well as AlphaFold3 with MSAs[33, 52]. The NDM-1 family was predicted with the substrate meropenem and two Zn ions, and predictions were compared to the experimentally solved complex 4EYL[39].

Pocket RMSD quantified the structural similarity of the ligand-binding pocket between a prediction and the reference structure 4EYL. Pocket residues were identified independently in each structure as any residue with an atom within 4.5 Å of any ligand atom (the two Zn^2+^ ions and the substrate); no residue correspondence was assumed between the reference and predicted pocket at this stage. C*α*-only coordinates for the resulting residue sets were extracted from each structure and superposed with USalign[53] in its default sequence-independent structural-alignment mode, which determines its own residue correspondence between the two sets rather than assuming a fixed pairing. RMSD_pocket_ is the RMSD reported for this alignment. Because the two pocket definitions are derived independently and USalign performs its own correspondence, RMSD_pocket_ reflects the geometric similarity of the two pockets as defined by each structure’s own ligand contacts, rather than a residue-matched deviation.

### 6.9. Candidate Selection and Filtering

#### 6.9.1. Metallo-β-lactamase

Candidates were ranked by a composite score with a required-contact gate applied first. Eight active-site residues were treated as required contacts: the Zn1-coordinating triad (H120, H122, H189), the Zn2-coordinating triad (D124, C208, H250), and the two substrate-binding residues (K211, N220)[39]. Letting *n*_missing_ be the number of these eight not in contact with their ligand in a given prediction (0–8), the composite score was

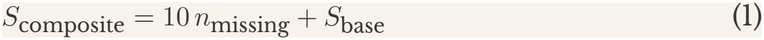

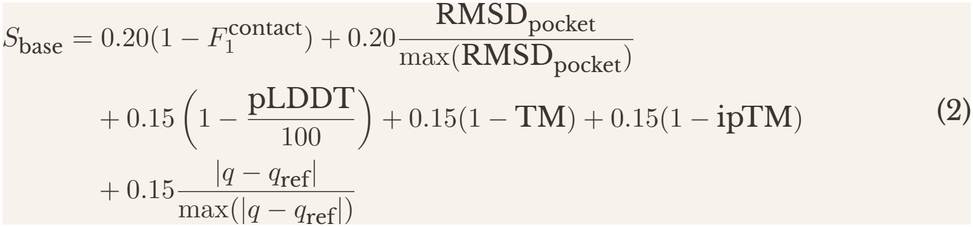

where 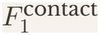 is the contact F1-score, RMSD_pocket_ is the pocket RMSD, pLDDT is the predicted local distance difference test score, TM is the TM-score, ipTM is the interface predicted TM-score, *q* is the net charge, and *q*_ref_ is the reference charge[54]. Lower *S*_composite_ indicates a better candidate. Because *S*_base_ is bounded by 1 and each missing required contact adds 10, a candidate missing any required contact is always ranked below every candidate missing fewer, regardless of its value on the other five metrics.

Candidates were additionally required to pass a set of physical-plausibility filters on the predicted binding pose: substrate-to-Zn1 distance *≤* 4.0 Å, substrate-to-Zn2 distance *≤* 4.0 Å, distance from the substrate’s metal-proximal atoms to the Zn–Zn bimetal midpoint *≤* 6.5 Å, and net charge *≤ –*5.0.

Top-ranked candidates passing all filters were manually inspected to confirm correct ligand binding geometry before selection for experimental validation.

#### Green fluorescent protein

Sequences generated under the three evaluated *α*_max_ conditions (8, 10, 12; see Sequence Generation) were pooled across runs. Where the same sequence occurred in more than one run, the copy from the run with the higher mean pLDDT was kept. Candidates were then filtered to net charge *≤ –*5 and at most three cysteines, and restricted to sequences with a PASS maturation verdict (Supplementary Methods) in one of the four colour classes targeted by conditioning: green, yellow, cyan/teal, and blue.

Within each colour class, the remaining candidates were ranked by a weighted score,

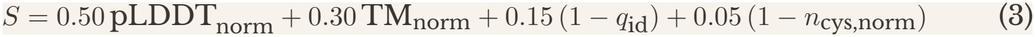

where pLDDT, TM-score, and cysteine count are min-max normalised across the full pool of PASS candidates, and *q*_id_ is the maximum sequence identity to any training-set sequence. Up to ten top-ranked candidates were retained per colour class (fewer where fewer than ten PASS candidates existed). The final two candidates per colour were selected from this ranked shortlist by manual review against AlphaFold3 pLDDT and pTM values and surface hydrophobicity, computed independently for the same sequences but not included in the composite score, giving a final set of eight candidates, two per colour class, all drawn from the *α*_max_ = 10 or *α*_max_ = 12 runs.

### 6.10. Surface redesign of NDM-1 candidate

Surface redesign was applied to the NDM-1 design with the best *in silico* metrics: NDM-D2. Its AlphaFold3 model, predicted in complex with the meropenem and two Zn ions, served as the structure reference. Surface positions were defined on the full complex: per-residue SASA was calculated with the Shrake-Rupley algorithm (500 sphere points; biotite[55]) and normalised to the maximum residue accessibility of Tien et al. [56] and residues with a relative accessibility *≥* 0.25 whose atoms were all more than 6 Å away from the meropenem and any Zn ion were designated designable surface positions. This identified 97 out of 255 residues. Sequences for the surface positions were sampled with LigandMPNN (weights `ligandmpnn_v_32_010_25`) as implemented in Foundry, with the backbone, meropenem, and Zn coordinates fixed as structural context[57]. Sampling was performed at a temperature of 0.2 with cysteine excluded at the designed positions with no other biases. Sixteen sequences were sampled and each redesigned sequence was then predicted with AlphaFold3[52], and the top-ranked models were compared by pLDDT (overall, protein, ligand and Zn), pTM, ipTM, *Cα* RMSD to the parent model and surface hydrophobicity. Surface hydrophobicity was assessed on the isolated protein chain as the fraction of exposed residues (relative accessibility *≥* 0.25) that were Ala, Val, Leu, Ile, Met, Phe, Pro or Trp. Hydrophobic patches were defined as sets of two or more such residues linked by *Cβ*-*Cβ* distances *≤* 8 Å(*Cα* for Gly), with patch area taken as the summed residue solvent-accessible surface area.

### 6.11. Experimental validation

#### 6.11.1. Cloning and Transformation of designed proteins

DNA fragments containing a BsaI recognition site at each end were commercially synthesized and cloned into the LM627 plasmid[58] using Golden Gate cloning[59]. The assembly reaction was incubated at 37 ^*≥*^C for 50 min, followed by a 10 min incubation at 60 ^*≥*^C to inactivate the enzymes. After assembly, the plasmids were transformed into E.coli Bl21 cells using heat shock at 42 ^*≥*^C. The transformed cells were recovered in 500 µl S.O.C. media for 1h. Transformants were streaked onto agar plates containing 50 µg/ml Kanamycin and 34 µg/ml Chloramphenicol and incubated overnight at 37 ^*≥*^C.

#### 6.11.2. Protein expression and purification

Single colonies were picked and grown overnight in 5 mL Terrific Broth (TB) medium supplemented with 50 µg/ml Kanamycin and 34 µg/ml Chloramphenicol. The following day 1% (v/v) of the overnight culture was transferred into 50 mL of TB containing 50 µg/ml Kan and 34 µg/ml Cam. Cells were grown at 37 ^*≥*^C until an *OD*_600_ of 0.5 was reached. Protein expression was induced by adding IPTG to a final concentration of 0.5 mM. To the NDM-1 desings, 100 uM of ZnCl were added into the expression cultures. The expression was left incubating for 18 hours at 20 ^*≥*^C with continuous shaking. Following expression, cultures were harvested by centrifugation at 4000 rpm for 20 min. The cell pellets were resuspended in 10 ml lysis buffer (20 mM HEPES, 150 mM NaCl, 20 mM imidazole). Dnase I (0.1 mg/ml), lysozyme (1 mg/mL), and protease inhibitor were added to the suspension. The cell suspensions were incubated on ice for 30 min, followed by three freeze thaw cycles. For each cycle, the samples were frozen at −80 ^*≥*^C followed by complete thawing in a water bath. The lysates were centrifuged to remove insoluble cellular material, and the resulting soluble lysate was collected for purification. Purification was carried out using immobilized metal affinity chromatography (IMAC). A 24-well filter plate containing 2 mL nickel affinity resin per well was equilibrated with binding buffer (20 mM HEPES, 150 mM NaCl, 20 mM imidazole). The soluble lysates were added to the beads and incubated for 30 min at 4 ^*≥*^C to allow protein binding. A vaccum manifold was applied to remove the lysate. The beads were washed with binding buffer, and the protein was eluted using 1 mL of elution buffer containing 500 mM imidazole. Elution was carried out sequentially in steps of 500 µl.

To further purify the protein, a Sephadex 75 increase 100/300GL column was used for size-exclusion (SEC). The proteins were eluted in a low-salt buffer containing 50 mM NaCl and 20 mM HEPES at pH 7.5. The resulting protein fractions were collected and concentrated to 500 µl using a spin column with a cutoff of 10 kDa.

#### 6.11.3. Protein characterization

To validate protein elution and purity, Western blotting was performed on the eluted samples collected after SEC. The membranes were incubated with an anti-His antibody to detect the His-tagged recombinant proteins. The proteins were visualized using chemiluminescent detection.

#### 6.11.4. Fluorescence spectroscopy

Fluorescence emission and excitation spectra of the purified fluorescent proteins were recorded in cuvettes using a spectrofluorometer. For emission spectra, samples were excited at the respective excitation wavelength, and fluorescence emission was recorded over the relevant wavelength range4. For excitation spectra, the samples were emitted at a fixed wavelength and the excitation wavelength was recorded over the relevant wavelength range4. A slit width of 5 nm was used for both excitation and emission, and spectra were recorded at a scan rate of 50 nm/min.

#### 6.11.5. Activity measurements

The enzymatic activity of NDM-1 designs was measured using a Nitrocefin-based assay. Each reaction had a final volume of 100 µL and contained 20 mM HEPES buffer, 10 µM *ZnCl*_2_, and 10 µM Nitrocefin. Wild-type NDM-1 (WT) was added at a final concentration of 50 nM. For each NDM-1 design 10 µl was added to the reaction mixture.

The hydrolysis of Nitrocefin was monitored by measuring absorbance at 490 nm over a period of 10 minutes. All reactions were blanked against a reference mixture containing the assay components but no enzyme. The initial linear rates of change in absorbance (Δ*A*_490_/min) were determined and normalized to protein concentration estimated by Western blot analysis. The normalized values were used to evaluate and compare the enzymatic activities of the different NDM-1 designs.

