## Supplementary text for "Conditional Dirichlet Flow Matching for exploring the sequence space of protein families"

### 7. Supplementary

#### Supplementary Results: Training and inference speeds

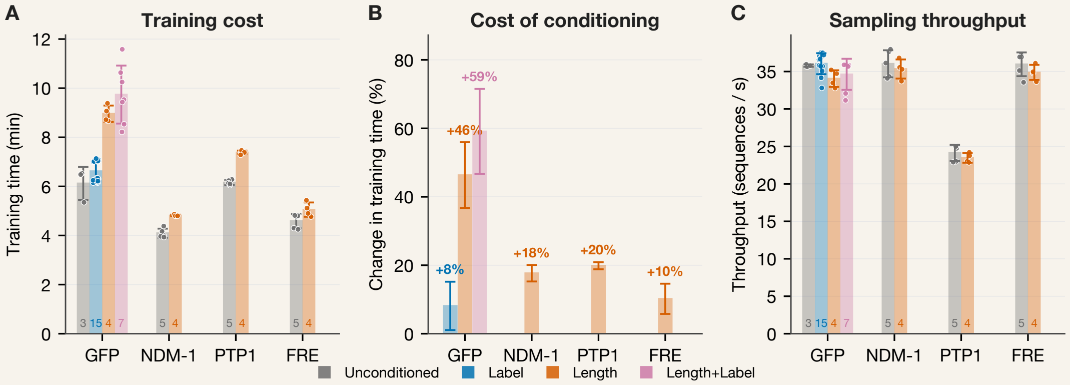


**Figure S1:** ***Conditioning raises training time but does not change sampling throughput.*** *All panels only contain data from single-GPU A100 runs.* ***(A)*** *Wall-clock training time per protein family, split by conditioning mode. Bars are means, error bars are one standard deviation, and each point is one run; run counts are given at the foot of each bar.* ***(B)*** *Training time of each conditioning mode as a percentage change from the unconditioned runs of the same family. Error bars propagate the standard error of both means through the ratio. Length conditioning costs 10–20% for FRE, NDM-1 and PTP1 and 46% for GFP; label conditioning costs 8 ± 7%, and the two together 59%.* ***(C)*** *Sustained sampling throughput per family from runs of at least 100 sequences. Bars are means, error bars are one standard deviation, and each point is one run. Throughput is set by the family. Within every family, conditioning changes it by less than 5%, comparable to the run-to-run spread: GFP, the only family run with all four modes, samples at 34–36 sequences/s under each of them. Across families, GFP, NDM-1 and FRE sample at 35–36 sequences/s and PTP1 at about 25 sequences/s. Sampling cost per sequence scales with the width of the alignment the model is defined on, which is 819 columns for PTP1 and 606–617 for the others. Runs of 100–300 sequences, which include most length-conditioned runs, read a few percent below the sustained rate because the fixed start-up cost is not fully amortised.*

#### Supplementary Results: $\alpha_{\max}$ comparison

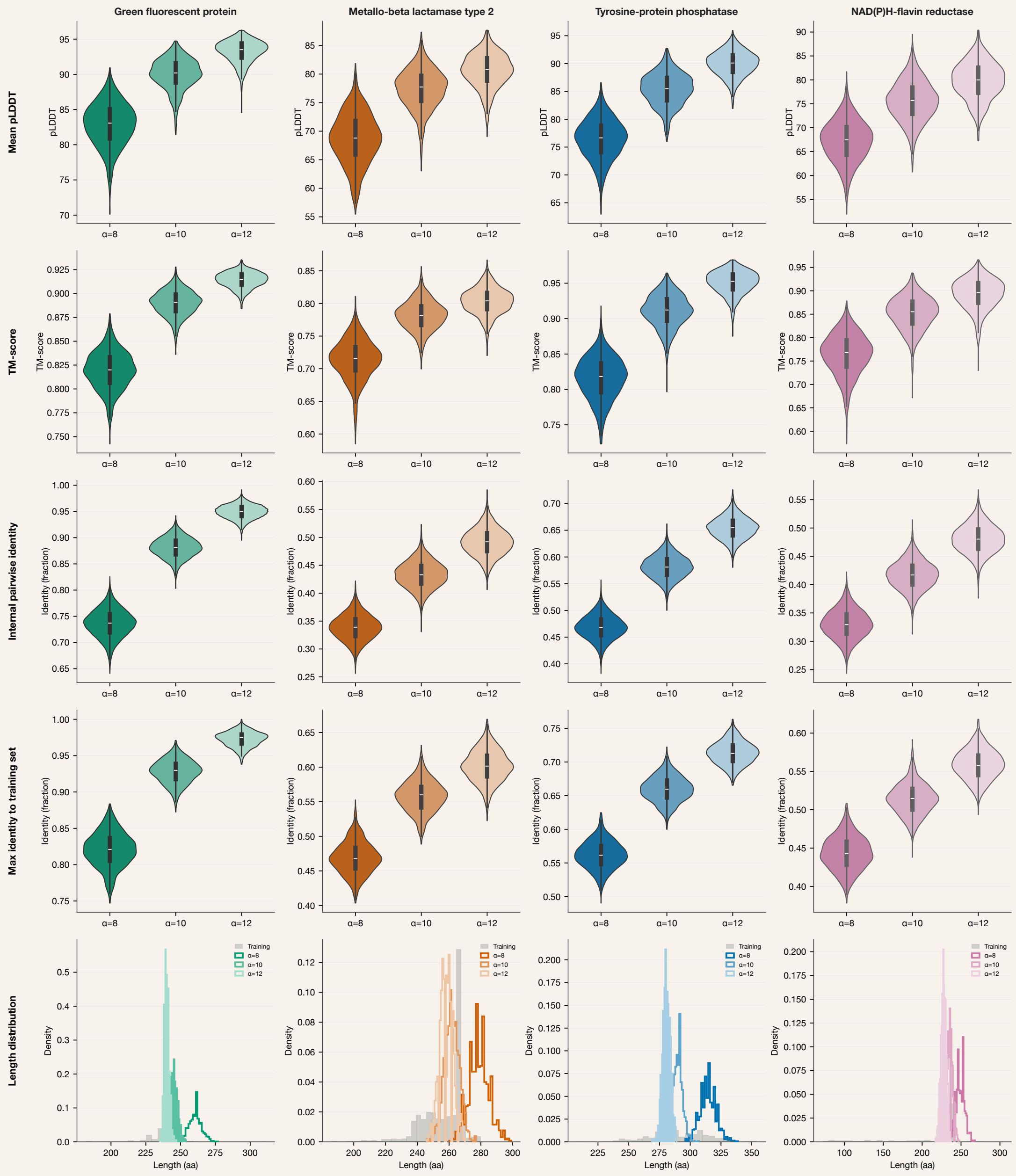


**Figure S2: *Higher $\alpha_{\max}$ raises structural confidence and identity to the training set, and narrows the length distribution.*** *Rows from top: mean pLDDT, TM-score, internal pairwise identity, mean maximum identity to the training set, length distribution. Violins show 1000 sequences per $\alpha_{\max}$ (Chai-1). The bottom row compares training (grey) with generated sequences.*

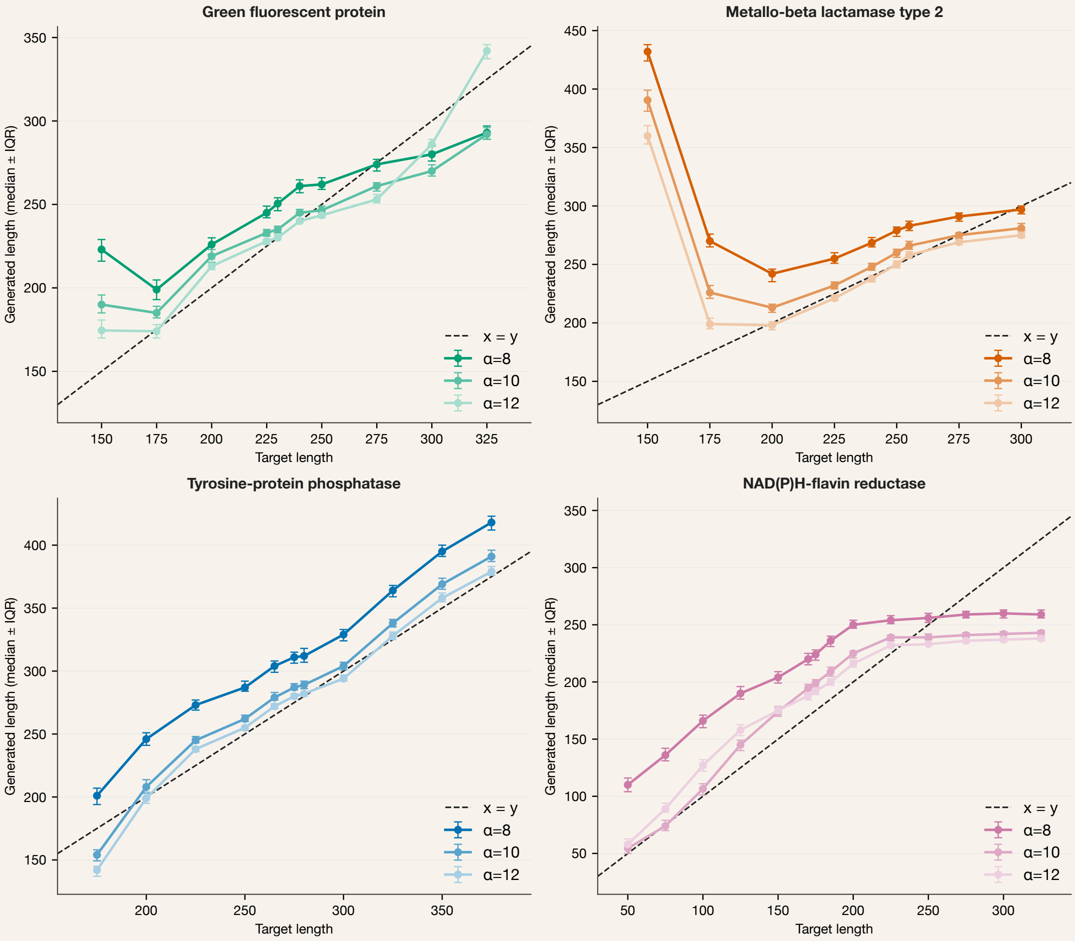


**Figure S3:** ***Length conditioning error falls with $\alpha_{\max}$ for targets with training support.*** *Median generated length (± IQR) against the requested length at $\alpha_{\max} = 8, 10, 12$. ($n = 150$ sequences per target length). The dashed line is perfect conditioning.*

**Table S1:** *Repetition in unconditioned GFP samples at different values of $\alpha_{\max}$ ($n = 1000$ per value). Identical to training is the percentage of generated sequences that are exact copies of a training sequence. Distinct training sequences copied is the number of different training sequences that these copies come from. Duplicated is the percentage of generated sequences that are exact copies of another generated sequence, calculated as one minus the number of unique sequences divided by 1000.*

| $\alpha_{\max}$ | Identical to training (%) | Distinct training sequences copied | Duplicated generated sequences (%) |
|---|---|---|---|
| 8 | 0.0 | 0 | 0.0 |
| 10 | 0.0 | 0 | 0.0 |
| 12 | 0.2 | 2 | 0.0 |
| 13 | 0.5 | 3 | 0.4 |
| 14 | 1.2 | 2 | 1.7 |
| 15 | 4.2 | 7 | 7.8 |

#### Supplementary Results: FoldSeek success rates

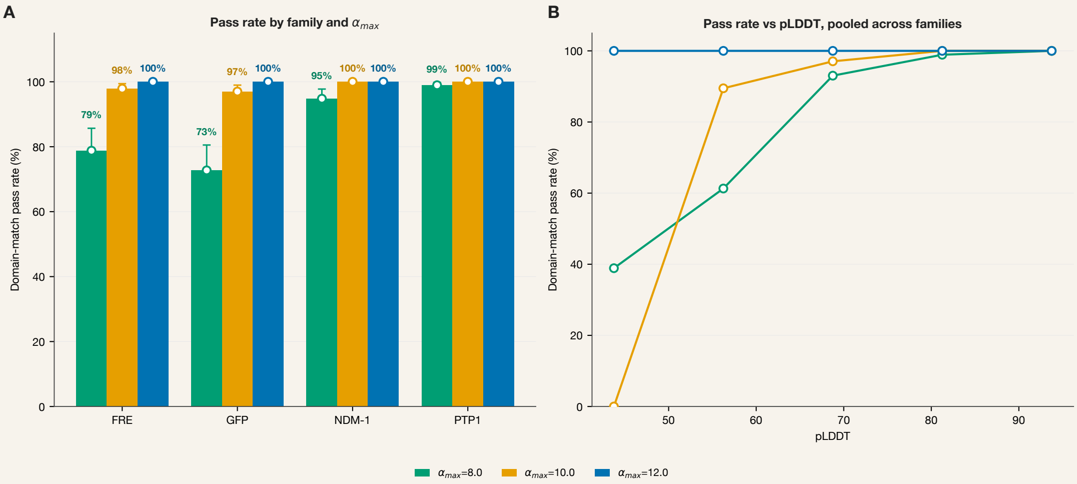


**Figure S4:** ***FoldSeek domain-match pass rate across sampling concentration ($\alpha_{\max}$) and structural confidence. (A)*** *Domain-match pass rate for each of the four evaluated protein families at $\alpha_{\max} = 8$, 10, and 12 ($n = 100$ sequences per condition). A sequence was scored as passing if at least one high-confidence FoldSeek hit shared a Pfam or InterPro identifier with the sequence's target family (see Methods). Error bars show Wilson 95% confidence intervals [60]. Pass rate increased with $\alpha_{\max}$ for every family, with the largest gains in GFP (P42212) and FRE (P43127), both of which rose from the lowest $\alpha_{\max} = 8$ baseline to ≥97% by $\alpha_{\max} = 10$.* ***(B)*** *Domain-match pass rate as a function of predicted structural confidence (pLDDT), binned and pooled across all four families, shown separately for each $\alpha_{\max}$. The low-pLDDT region of the $\alpha_{\max} \geq 10$ curves is sparsely sampled, drawn primarily from a small number of GFP sequences, and should be weighted accordingly.*

#### Supplementary Results: Co-conditioning

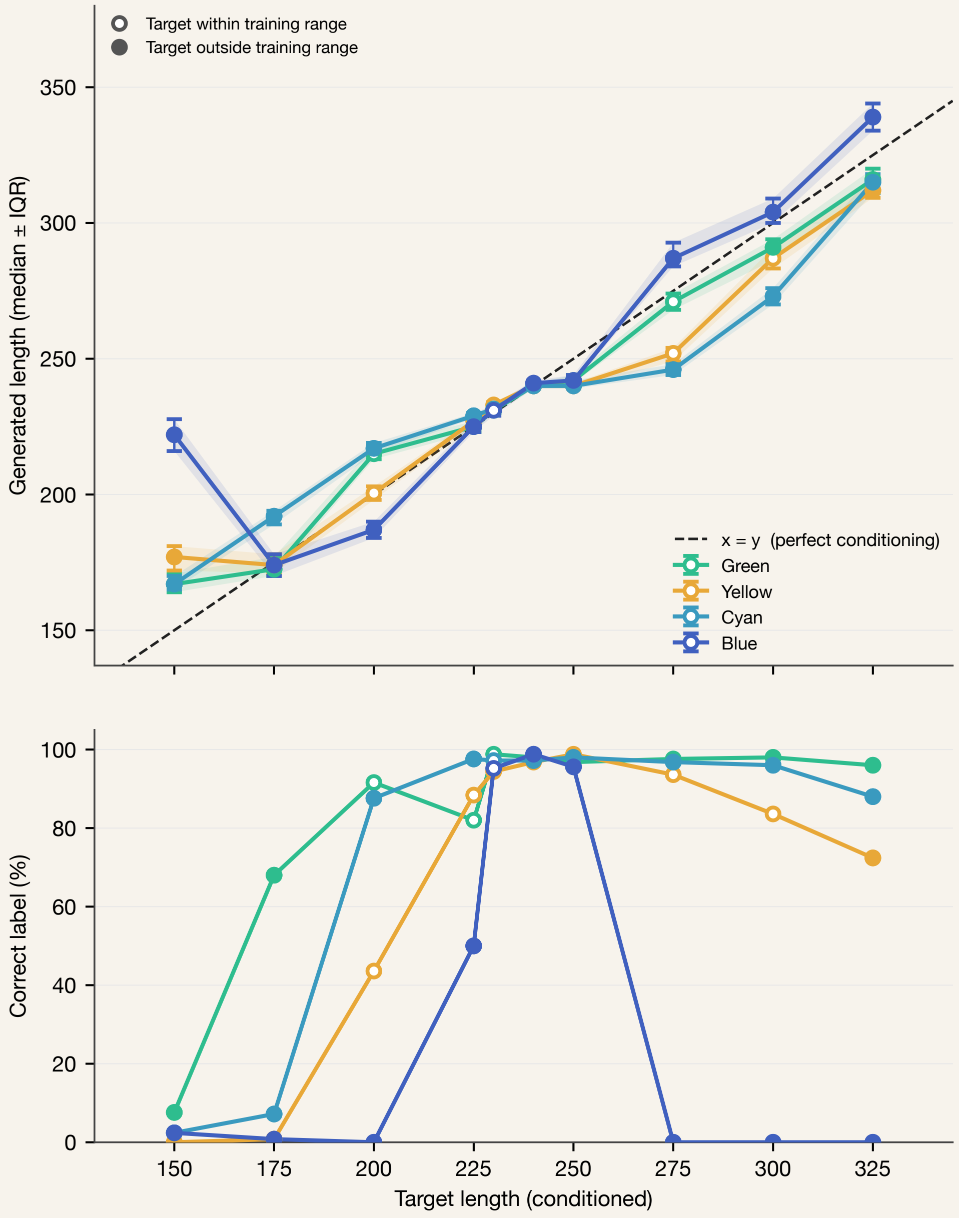


**Figure S5:** ***Co-conditioning keeps the length pattern of length-only conditioning, with label accuracy reflecting the necessity of in-distribution training examples.*** *Top: median generated length (± IQR) against the target length, per requested label, at* $\alpha_{\max} = 12$*. Bottom: percentage of sequences with the requested label at each target. Open markers: target within the training length range; filled markers: outside it.*

#### Supplementary Results: Experimental data

**Table S2:** *Excitation and emission peaks of the GFP designs.*

| Protein | Emission peak (nm) | Excitation peak (nm) |
|---|---|---|
| GFP WT | 507.5 | 393.5 |
| GFP-1 | 507.7 | 392.5 |
| GFP-2 | 506.5 | 393.0 |
| YFP-1 | 507.5 | 394.5 |
| CFP-1 | 506.0 | 393.5 |
| BFP-1 | 446.5 | 378.5 |
| BFP-2 | – | – |
| Extended GFP-1 | 507.5 | 394.5 |

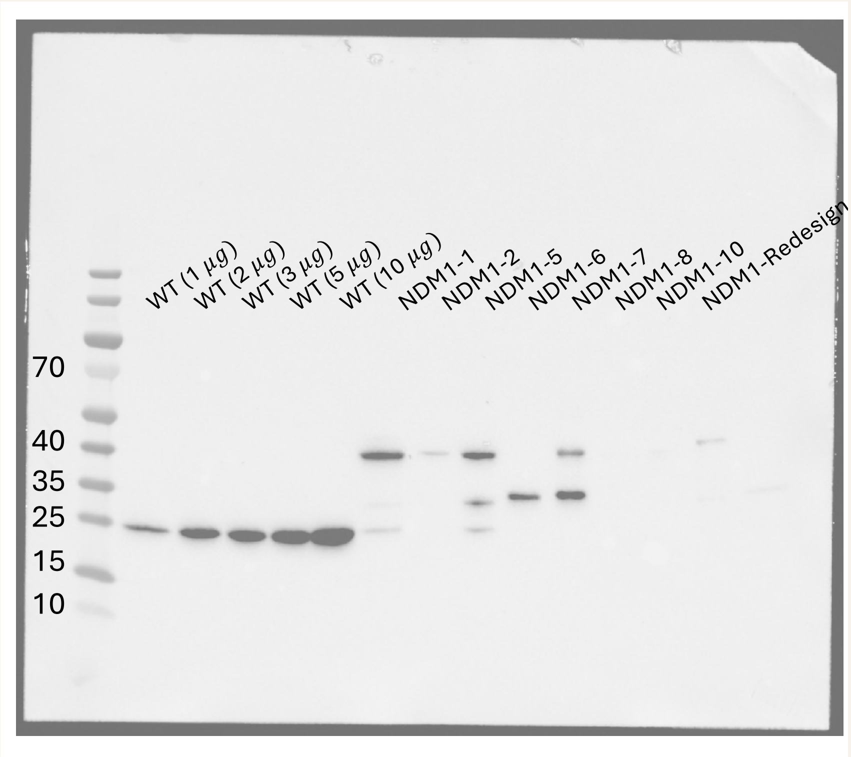


**Figure S6:** ***Western Blot of NDM1-WT in different concentrations and NDM-1 Designs.***

#### Supplementary Methods: Definition of conditioning success

Generated sequences were evaluated at chromophore-region positions to assign (i) a maturation verdict (PASS/WARN/FAIL) and (ii) a colour class, which was compared against the requested conditioning label to score conditioning accuracy, reported in Table S3.

**Table S3:** *Positions defining verdict and colour class.*

| Pos | WT | Accepted residues | Ref |
|---|---|---|---|
| 67 | G | G only (else FAIL) | [61] |
| 96 | R | R, K; A rescued if paired with 183R | [62, 63] |
| 183 | Q | Q, R (R is the 96A rescue) | [63] |
| 65 | S | S, T, A, G, C, L | [64] |
| 66 | Y | Y (green/yellow), W (cyan), H (blue) | [64] |
| 203 | T | T, Y, H, C (splits yellow from green within Y66) | [65] |

The verdict logic is as follows: FAIL if position 67 is not glycine, or if position 96 is non-catalytic and not rescued by the 96A/183R pair. All substitutions listed at 65 and 66 above are PASS-compatible; WARN is reserved for residues at 65 or 66 falling outside this set, which did not occur among the reported sequences. Colour class is assigned from position 66, with position 203 resolving the green/yellow distinction when 66 = Y.

**Table S4:** *Catalytic residues of the four protein families.*

| Protein | Residue and position | Task | Ref |
|---|---|---|---|
| NDM-1 a structural view papers | H120 | $Zn^{2+}$(I) coordinating residue | [17, 66, 67] crystal structure |
| | H122 | $Zn^{2+}$(I) coordinating residue | [17, 66, 67] |
| | D124 | $Zn^{2+}$(II) coordinating residue | [17, 66, 67] |
| | H189 | $Zn^{2+}$(I) coordinating residue | [17, 66, 67] |
| | C208 | $Zn^{2+}$(II) coordinating residue | [17] |
| | N220 | Substrate binding through hydrogen bond | [17] |
| | H250 | $Zn^{2+}$(II) coordinating residue | [17, 66, 67] |
